# A translation control module coordinates germline stem cell differentiation with ribosome biogenesis during *Drosophila* oogenesis

**DOI:** 10.1101/2021.04.04.438367

**Authors:** Elliot T. Martin, Patrick Blatt, Elaine Ngyuen, Roni Lahr, Sangeetha Selvam, Hyun Ah M. Yoon, Tyler Pocchiari, Shamsi Emtenani, Daria E. Siekhaus, Andrea Berman, Gabriele Fuchs, Prashanth Rangan

## Abstract

Ribosomal defects perturb stem cell differentiation, causing diseases called ribosomopathies. How ribosome levels control stem cell differentiation is not fully known. Here, we discovered three RNA helicases are required for ribosome biogenesis and for *Drosophila* oogenesis. Loss of these helicases, which we named Aramis, Athos and Porthos, lead to aberrant stabilization of p53, cell cycle arrest and stalled GSC differentiation. Unexpectedly, Aramis is required for efficient translation of a cohort of mRNAs containing a 5’-Terminal-Oligo-Pyrimidine (TOP)-motif, including mRNAs that encode ribosomal proteins and a conserved p53 inhibitor, <u>No</u>vel <u>N</u>ucleolar protein 1 (Non1). The TOP-motif co-regulates the translation of growth-related mRNAs in mammals. As in mammals, the La-related protein co-regulates the translation of TOP-motif containing RNAs during *Drosophila* oogenesis. Thus, a previously unappreciated TOP-motif in *Drosophila* responds to reduced ribosome biogenesis to co-regulate the translation of ribosomal proteins and a p53 repressor, thus coupling ribosome biogenesis to GSC differentiation.

## Introduction

All life depends on the ability of ribosomes to translate mRNAs into proteins. Despite this universal requirement, ribosome biogenesis is not universally equivalent. Stem cells, the unique cell type that underlies the generation and expansion of tissues, in particular have a distinct ribosomal requirement (Gabut et al., 2020; Sanchez et al., 2016; Woolnough et al., 2016; Zahradkal et al., 1991; Zhang et al., 2014). Ribosome production and levels are dynamically regulated to maintain higher amounts in stem cells (Fichelson et al., 2009; Gabut et al., 2020; Sanchez et al., 2016; Woolnough et al., 2016; Zahradkal et al., 1991; Zhang et al., 2014). For example, ribosome biogenesis components are often differentially expressed, as observed during differentiation of embryonic stem cells, osteoblasts, and myotubes (Gabut et al., 2020; Watanabe-Susaki et al., 2014; Zahradkal et al., 1991). In some cases, such as during *Drosophila* germline stem cell (GSC) division, ribosome biogenesis factors asymmetrically segregate during asymmetric cell division, such that a higher pool of ribosome biogenesis factors is maintained in the stem cell compared to the daughter cell (Blatt et al., 2020a; Fichelson et al., 2009; Zhang et al., 2014). Reduction of ribosome levels in several stem cells systems can cause differentiation defects (Corsini et al., 2018; Fortier et al., 2015; Khajuria et al., 2018; Zhang et al., 2014). In *Drosophila,* perturbations that reduce ribosome levels in the GSCs result in differentiation defects causing infertility (Sanchez et al., 2016). Similarly, humans with reduced ribosome levels are afflicted with clinically distinct diseases known as ribosomopathies, such as Diamond-Blackfan anemia, that often result from loss of proper differentiation of tissue-specific progenitor cells (Armistead and Triggs-Raine, 2014; Barlow et al., 2010; Brooks et al., 2014; Higa-Nakamine et al., 2012; Lipton et al., 1986; Mills and Green, 2017). However, the mechanisms by which ribosome biogenesis is coupled to proper stem cell differentiation remain incompletely understood.

Ribosome production requires the transcription of ribosomal RNAs (rRNAs) and of mRNAs encoding ribosomal proteins (Bousquet-Antonelli et al., 2000; de la Cruz et al., 2015; Granneman et al., 2011, 2006; Tafforeau et al., 2013; Venema et al., 1997). Several factors, such as helicases and endonucleases, transiently associate with maturing rRNAs to facilitate rRNA processing, modification, and folding (Granneman et al., 2011; Sloan et al., 2017; Tafforeau et al., 2013; Watkins and Bohnsack, 2012). Ribosomal proteins are imported into the nucleus, where they assemble with rRNA to form the small 40S and large 60S ribosome subunits, which are then exported to the cytoplasm (Baxter-Roshek et al., 2007; Decatur and Fournier, 2002; Granneman et al., 2011, 2006; Koš and Tollervey, 2010; Nerurkar et al., 2015; Tafforeau et al., 2013; Zemp and Kutay, 2007). Loss of RNA Polymerase I transcription factors, helicases, exonucleases, large or small subunit ribosomal proteins, or other processing factors all compromise ribosome biogenesis and trigger diverse stem cell-related phenotypes (Brooks et al., 2014; Calo et al., 2018; Mills and Green, 2017; Sanchez et al., 2016; Yelick and Trainor, 2015; Zhang et al., 2014).

Nutrient availability influences the demand for *de novo* protein synthesis and thus ribosome biogenesis (Anthony et al., 2000; Hong et al., 2012; Mayer and Grummt, 2006; Shu et al., 2020). In mammals, nearly all of the mRNAs that encode the ribosomal proteins contain a Terminal Oligo Pyrimidine (TOP) motif within their 5’ untranslated region (UTR), which regulates their translation in response to nutrient levels (Fonseca et al., 2015; Hong et al., 2017; Lahr et al., 2017; Tcherkezian et al., 2014). Under growth-limiting conditions, La related protein 1 (Larp1) binds to the TOP sequences and to mRNA caps to inhibit translation of ribosomal proteins (Fonseca et al., 2015; Jia et al., 2021; Lahr et al., 2017; Philippe et al., 2018). When growth conditions are suitable, Larp1 is phosphorylated by the nutrient/redox/energy sensor mammalian Target of rapamycin (mTOR) complex 1 (mTORC1), and does not efficiently bind the TOP sequence, thus allowing for efficient translation of ribosomal proteins (Fonseca et al., 2018, 2015; Hong et al., 2017; Jia et al., 2021). In some instances, Larp1 binding can also stabilize TOP-containing mRNAs (Aoki et al., 2013; Berman et al., 2020; Gentilella et al., 2017; Ogami et al., 2020), linking mRNA translation with mRNA stability to promote ribosome biogenesis (Aoki et al., 2013; Berman et al., 2020; Fonseca et al., 2018, 2015; Hong et al., 2017; Lahr et al., 2017; Ogami et al., 2020; Philippe et al., 2018). Cellular nutrient levels are known to affect stem cell differentiation and oogenesis in *Drosophila* (Hsu et al., 2008), however whether TOP motifs exist in *Drosophila* to coordinate ribosome protein synthesis is unclear. The *Drosophila* ortholog of Larp1, La related protein (Larp) is required for proper cytokinesis and meiosis in *Drosophila* testis as well as for female fertility, but its targets remain undetermined (Blagden et al., 2009; Ichihara et al., 2007).

Germline depletion of ribosome biogenesis factors manifests as a stereotypical GSC differentiation defect during *Drosophila* oogenesis (Sanchez et al., 2016). Female *Drosophila* maintain 2-3 GSCs in the germarium (**Figure 1A**) (Kai et al., 2005; Twombly et al., 1996; Xie, 2000; Xie and Li, 2007; Xie and Spradling, 1998). Asymmetric cell division of GSCs produces a self-renewing daughter GSC, and a differentiating daughter, called the cystoblast (CB) (Chen and McKearin, 2003; McKearin and Ohlstein, 1995). This asymmetric division is unusual: following mitosis, the abscission of the GSC and CB is not completed until the following G2 phase (**Figure 1A’**) (De Cuevas and Spradling, 1998; Hsu et al., 2008). The GSC is marked by a round structure called the spectrosome, which elongates and eventually bridges the GSC and CB, similar to the fusomes that connect differentiated cysts (**Figure 1A’**). During abscission the extended spectrosome structure is severed and a round spectrosome is established in the GSC and the CB (De Cuevas and Spradling, 1998; Hsu et al., 2008). Ribosome biogenesis defects result in failed GSC-CB abscission, causing cells to accumulate as interconnected cysts marked by a fusome-like structure called “stem cysts” (**Figure 1A’**) (Mathieu et al., 2013; Sanchez et al., 2016). In contrast with differentiated cysts (McKearin and Ohlstein, 1995; McKearin and Spradling, 1990; Ohlstein and McKearin, 1997), these stem cysts lack expression of the differentiation factor Bag of Marbles (Bam), do not differentiate, and typically die, resulting in sterility (Sanchez et al., 2016). How proper ribosome biogenesis promotes GSC abscission and differentiation is not known.

**Figure 1:**
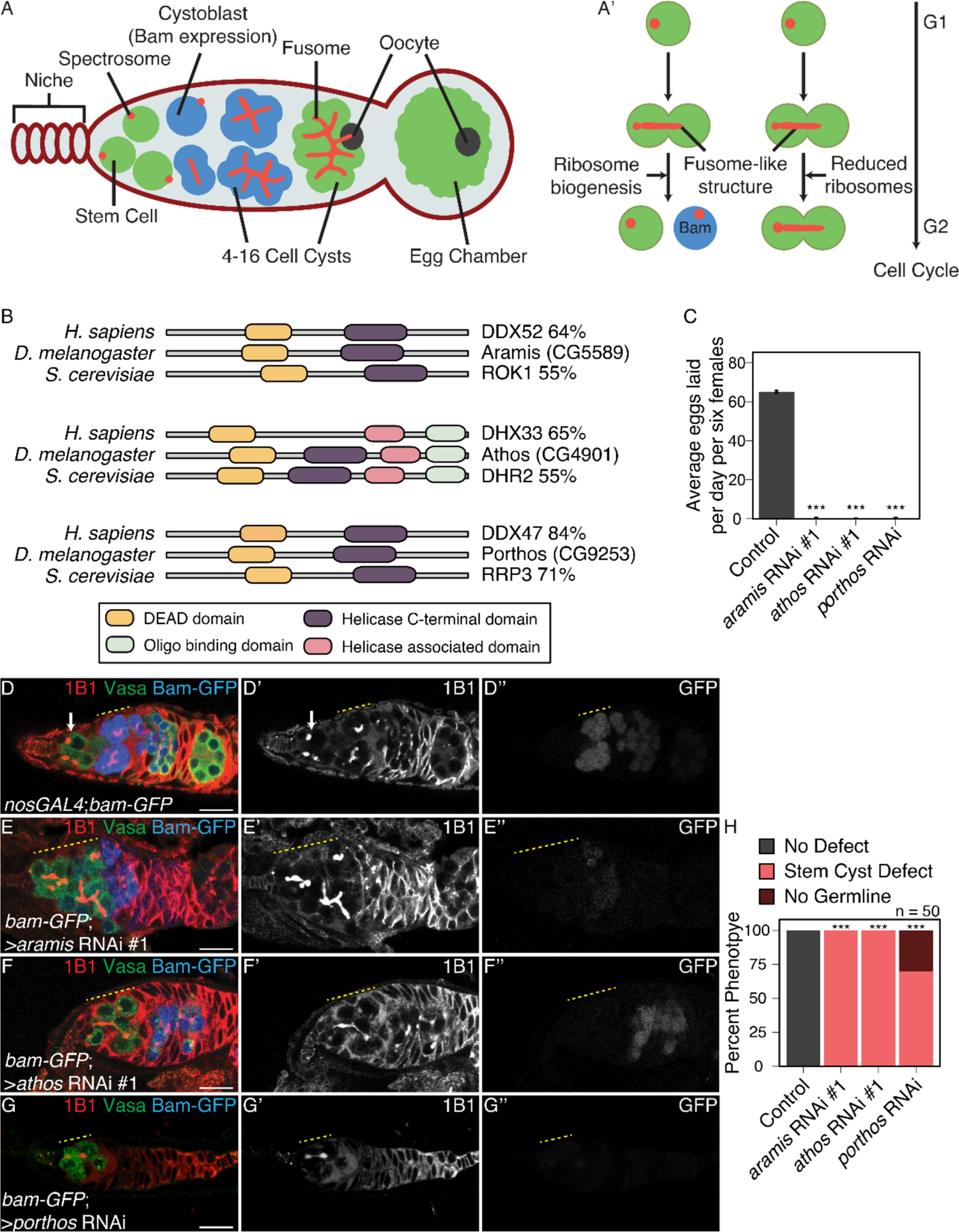
RNA helicases Aramis, Athos and Porthos are required for GSC differentiation. (**A**) Schematic of *Drosophila* germarium. Germline stem cells are attached to the somatic niche (dark red). The stem cells divide and give rise to a stem cell and a cystoblast (CB) that expresses the differentiation factor Bag-of-marbles (Bam). GSCs and CBs are marked by spectrosomes. The CB undergoes four incomplete mitotic divisions giving rise to a 16-cell cyst (blue). Cysts are marked by branched spectrosome structures known as fusomes (red). One cell of the 16-cell cyst is specified as the oocyte. The 16-cell cyst is encapsulated by the surrounding somatic cells giving rise to an egg chamber. (**A’**) Ribosome biogenesis promotes GSC cytokinesis and differentiation. Disruption of ribosome biogenesis results in undifferentiated stem cyst accumulation. (**B**) Representation of conserved protein domains for three RNA helicases in *Drosophila* compared to *H. sapiens* and *S. cerevisiae* orthologs. Percentage values represent similarity to *Drosophila* orthologs. (**C**) Egg laying assay after germline RNAi knockdown of *aramis*, *athos* or *porthos* indicating a loss of fertility compared to *nosGAL4*, driver control (n=3 trials). *** = p < 0.001, Tukey’s post-hoc test after one-way ANOVA, p < 0.001. Error bars represent standard error (SE). (**D-G’’**) Confocal micrographs of ovaries from control, *UAS-Dcr2;nosGAL4;bam-GFP* (**D-D’’**) and germline RNAi depletion targeting (**E-E’’**) *aramis*, (**F-F’’**) *athos* or (**G-G’’**) *porthos* stained for 1B1 (red, left grayscale), Vasa (green), and Bam-GFP (blue, right grayscale). Depletion of these genes (**E-G’’**) results in a characteristic phenotype in which early germ cells are connected marked by a 1B1 positive, fusome-like structure highlighted by a yellow dotted line in contrast to the single cells present in (**D-D’’**) controls (white arrow) or differentiating cysts (yellow dashed line). Bam expression, if present, is followed by loss of the germline. (**H**) Phenotype quantification of ovaries depleted of *aramis*, *athos* or *porthos* compared to control ovaries (n=50 ovarioles, df=2, *** = p < 0.001, Fisher’s exact tests with Holm-Bonferroni correction). Scale bars are 15 micron.

By characterizing three RNA helicases that promote ribosome biogenesis, we identified a translational control module that is sensitive to proper ribosome biogenesis and coordinates ribosome levels with GSC differentiation. When ribosome biogenesis is optimal, ribosomal proteins and a p53 repressor are both efficiently translated allowing for proper GSC cell cycle progression and its differentiation. However, when ribosome biogenesis is perturbed, we observe diminished translation of both ribosomal proteins and the p53 repressor. As a consequence, p53 is stabilized, cell cycle progression is blocked and GSC differentiation is stalled. Thus, our work reveals an elegant tuning mechanism that links ribosome biogenesis with a cell cycle progression checkpoint and thus stem cell differentiation. Given that ribosome biogenesis defects in humans result in ribosomopathies, which often result from stem cell differentiation defects, our data lay the foundation for understanding the etiology of developmental defects that arise due to ribosomopathies.

## Results

### Three conserved RNA helicases are required in the germline for GSC differentiation

We performed a screen to identify RNA helicases that are required for female fertility in *Drosophila*, and identified three predicted RNA helicases with previously uncharacterized functions, *CG5589*, *CG4901,* and *CG9253* (**Figure 1B-C**) (**Supplemental Table 1**) (Blatt et al., 2020b). We named these candidate genes *aramis*, *athos*, and *porthos,* respectively, after Alexandre Dumas’ three musketeers who fought in service of their queen. To further investigate how these helicases promote fertility, we depleted *aramis*, *athos*, and *porthos* in the germline using the germline-driver *nanos*-*GAL4* (*nosGAL4*) in combination with RNAi lines. We detected the germline and spectrosomes/fusomes in ovaries by immunostaining for Vasa and 1B1, respectively. In contrast to controls, *aramis*, *athos*, and *porthos* germline RNAi flies lacked spectrosome-containing cells, and instead displayed cells with fusome-like structures proximal to the self-renewal niche (**Figure 1D-H; Figure S1A-A’’’**). The cells in this cyst-like structure contained ring canals, a marker of cytoplasmic bridges, suggesting that they are indeed interconnected (**Figure S1B-B’’’**) (Zhang et al., 2014). In addition to forming cysts in an aberrant location, the *aramis*, *athos*, and *porthos* germline RNAi ovaries failed to form egg chambers (**Figure 1D-H**).

Aberrant cyst formation proximal to the niche could reflect stem cysts with GSCs that divide to give rise to CBs but fail to undergo cytokinesis or differentiated cysts that initiate differentiation but cannot progress further to form egg chambers. To discern between these possibilities, first we examined the expression of a marker of GSCs, phosphorylated Mothers against decapentaplegic (pMad). We observed pMad expression in the cells closest to the niche, but not elsewhere in the germline cysts of *aramis*, *athos*, and *porthos* germline RNAi flies (**Figure S1C-F’)** (Kai and Spradling, 2003). Additionally, none of the cells connected to the GSCs in *aramis*, *athos*, and *porthos* germline RNAi flies expressed the differentiation reporter *bamGFP* (**Figure 1D-G’’)** (McKearin and Ohlstein, 1995). Thus, loss of *aramis*, *athos*, or *porthos* in the germline results in the formation of stem cysts, however with variable severity. This variability could be due to a differential requirement for these genes or different RNAi efficiencies. Overall, we infer that Aramis, Athos, and Porthos are required for proper GSC cytokinesis to produce a stem cell and differentiating daughter.

### Athos, Aramis, and Porthos are required for ribosome biogenesis

We found that Aramis, Athos, and Porthos are conserved from yeast to humans (**Figure 1B**). The closest orthologs of Aramis, Athos, and Porthos are Rok1, Dhr2, and Rrp3 in yeast and DExD-Box Helicase 52 (DDX52), DEAH-Box Helicase 33 (DHX33), and DEAD-Box Helicase 47 (DDX47) in humans, respectively (Hu et al., 2011). Both the yeast and human orthologs have been implicated in rRNA biogenesis (Bohnsack et al., 2008; Khoshnevis et al., 2016; Martin et al., 2014; O’day et al., 1996; Sekiguchi et al., 2006; Tafforeau et al., 2013; Venema et al., 1997; Venema and Tollervey, 1995; Vincent et al., 2017; Zhang et al., 2011). In addition, the GSC-cytokinesis defect that we observed in *aramis*, *athos*, and *porthos* RNAi flies is a hallmark of reduced ribosome biogenesis in the germline (Sanchez et al., 2016). Based on these observations, we hypothesized that Aramis, Athos, and Porthos could enhance ribosome biogenesis to promote proper GSC differentiation.

Many factors involved in rRNA biogenesis localize to the nucleolus and interact with rRNA (Arabi et al., 2005; Grandori et al., 2005; Henras et al., 2008; Karpen et al., 1988). To detect the subcellular localization of Aramis and Athos, we used available lines that express Aramis::GFP::FLAG or Athos::GFP::FLAG fusion proteins under endogenous control. For Porthos, we expressed a Porthos::FLAG::HA fusion under the control of UASt promoter in the germline using a previously described approach (DeLuca and Spradling, 2018). We found that in the germline, Aramis, Athos and Porthos colocalized with Fibrillarin, which marks the nucleolus, the site of rRNA synthesis (**Figure 2A-C’’’**) (Ochs et al., 1985). Aramis was also in the cytoplasm of the germline and somatic cells of the gonad. To determine if Aramis, Athos, and Porthos directly interact with rRNA, we performed immunoprecipitation (IP) followed by RNA-seq. We found that rRNA immunopurified with Aramis, Athos, and Porthos (**Figure 2D-D’’, Figure S2A-A’’).** Thus, Aramis, Athos, and Porthos are present in the nucleolus and interact with rRNA, suggesting that they might regulate rRNA biogenesis.

**Figure 2.**
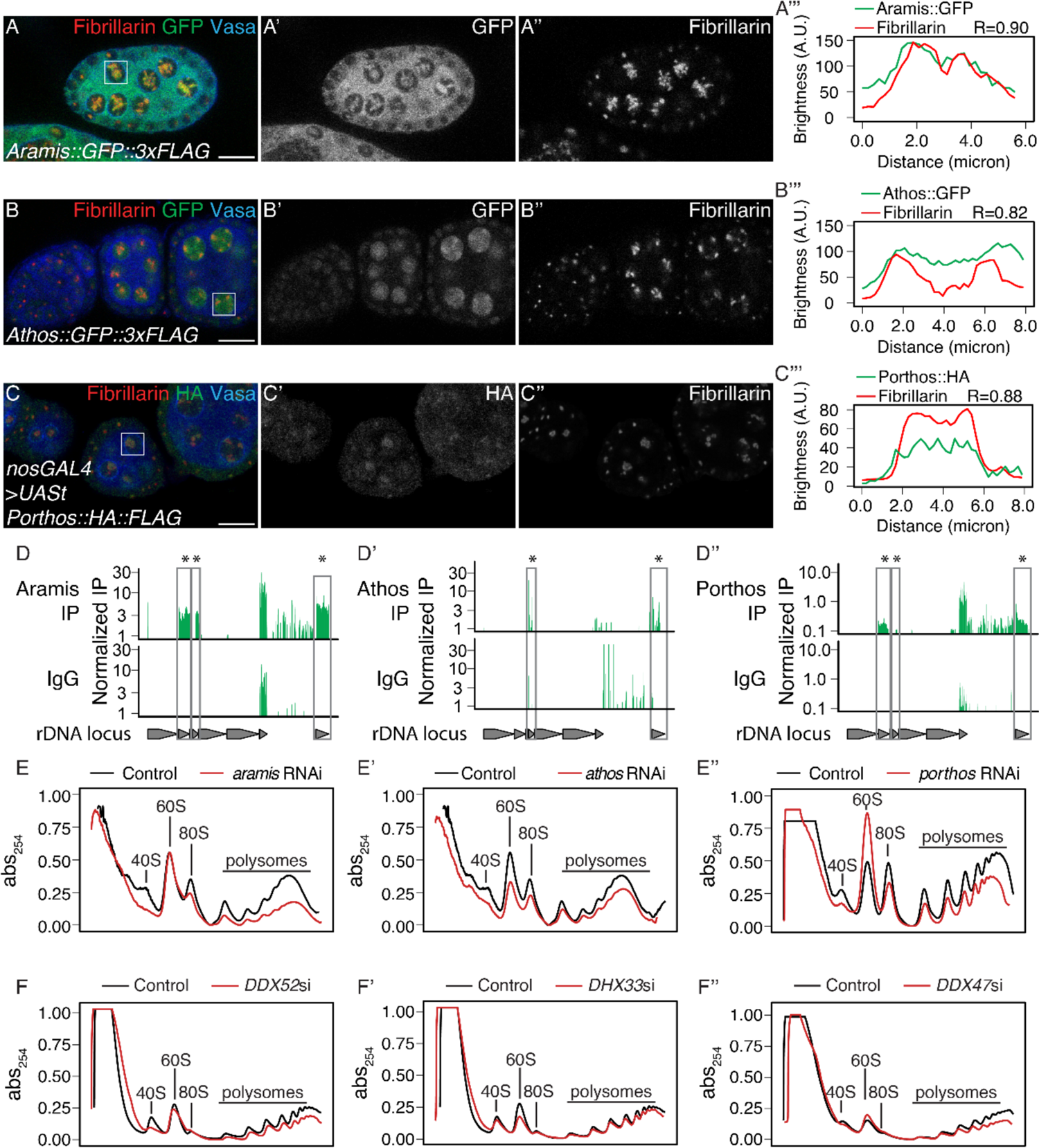
Athos, Aramis, and Porthos are required for efficient ribosome biogenesis. (**A-C’’**) Confocal images of ovariole immunostained for Fibrillarin (red, right grayscale), Vasa (blue), (**A-A’’**) Aramis::GFP, (**B-B’’**) Athos::GFP and (**C-C’’**) Porthos::HA (green, left grayscale). (**A’’’-C’’’**) Fluorescence intensity plot generated from a box of averaged pixels centered around the punctate of Fibrillarin in the white box. R values denote Spearman correlation coefficients between GFP and Fibrillarin from plot profiles generated using Fiji, taken from the nucleolus denoted by the white box. Aramis, Athos and Porthos are expressed throughout oogenesis and localize to the nucleolus. Aramis is also present in the cytoplasm. (**D-D’’**) RNA IP-seq of (**D**) Aramis, (**D’**) Athos, and (**D’’**) Porthos aligned to rDNA displayed as genome browser tracks. Bar height represents log scaled rRNA reads mapping to rDNA normalized to input and spike-in. Grey boxes outline rRNA precursors that are significantly enriched in the IP compared to the IgG control (bootstrapped paired t-tests, n=3, * = p-value < 0.05). (**E-E’’**) Polysome traces from *Drosophila* S2 cells treated with dsRNA targeting (**E**) *aramis*, (**E’**) *athos*, (**E’’**) *porthos* (red line) compared to a mock control (black line). *aramis* and *porthos* are required to maintain a proper 40S/60S ribosomal subunit ratio compared to control and have a smaller 40S/60S ratio. *athos* is required to maintain a proper 40S/60S ribosomal subunit ratio compared to control and has a larger 40S/60S ratio. Additionally, *aramis*, *athos*, and *porthos* are required to maintain polysome levels. (**F-F’’**) Polysome preparations from HeLa cells depleted of *DDX52*, *DHX33*, *DDX47*, and control siRNA treated cells. *DDX52*, *DHX33*, and *DDX47* are required to maintain a proper 40S/60S ribosomal subunit ratio. Additionally, all three are required to maintain polysome levels. Scale bar for all images is 15 micron.

Nucleolar size, and in particular nucleolar hypotrophy, is associated with reduced ribosome biogenesis and nucleolar stress (Neumüller et al., 2008; Zhang et al., 2011). If Aramis, Athos, and Porthos promote ribosome biogenesis, then their loss would be expected to cause nucleolar stress and a reduction in mature ribosomes. Indeed, immunostaining for Fibrillarin revealed hypotrophy of the nucleolus in *aramis, athos,* and *porthos* germline RNAi flies compared to in control flies, consistent with nucleolar stress (**Figure S2B-C**). Next, we used polysome profile analysis to evaluate the ribosomal subunit ratio and translation status of ribosomes in S2 cells depleted of *aramis*, *athos*, or *porthos* (Boamah et al., 2012; Õunap et al., 2013). We found that upon the depletion of all three helicases, the heights of the polysome peaks were reduced (**Figure 2E-E’’**). We found that depletion of *aramis* and *porthos* diminished the height of the 40S subunit peak compared to the 60S subunit peak, characteristic of defective 40S ribosomal subunit biogenesis (**Figure 2E, E’’**, **Figure S2D)** (Cheng et al., 2019), whereas *athos* depletion diminished the height of the 60S subunit peak compared to the 80S peaks, characteristic of a 60S ribosomal subunit biogenesis defect (**Figure 2E’, Figure S2D’**) (Cheng et al., 2019). RNAi-mediated depletion of the orthologs of these helicases in HeLa cells similarly affected the polysome profiles (**Figure 2F’-F’’, Figure S2E-G**). Taken together our findings indicate that these helicases promote ribosome biogenesis in *Drosophila* and mammalian cells.

### Aramis promotes cell cycle progression via p53 repression

Our data so far indicate that Aramis, Athos and Porthos promote ribosome biogenesis, which is known to be required for GSC abscission (Sanchez et al., 2016). Yet the connections between ribosome biogenesis and GSC abscission are poorly understood. To explore the connection, we further examined the *aramis* germline RNAi line, as its defect was highly penetrant but maintained sufficient germline for analysis (**Figure 1E, H**). First, we compared the mRNA profiles of *aramis* germline RNAi ovaries to *bam* germline RNAi to determine if genes that are known to be involved in GSC abscission have altered expression. We used germline *bam* depletion as a control because it leads to the accumulation of CBs with no abscission defects (Flora et al., 2018a; Gilboa et al., 2003; McKearin and Ohlstein, 1995; Ohlstein and McKearin, 1997), whereas loss of *aramis* resulted in accumulation of CBs that do not abscise from the GSCs.

We performed RNA-seq and found that 607 RNAs were downregulated and 673 RNAs were upregulated in *aramis* germline RNAi versus *bam* germline RNAi (cut-offs for differential gene expression were log_2_(foldchange) >|1.5|, FDR < 0.05) (**Figure S3A, Supplemental Table 2**). Gene Ontology (GO) analysis for biological processes on these genes encoding these differentially expressed mRNAs (Thomas et al., 2003) revealed that the genes that were downregulated upon *aramis* germline depletion were enriched for GO terms related to the cell cycle, whereas the upregulated genes were enriched for GO terms related to stress response (**Figure 3A, Figure S3B**). The downregulated genes included *Cyclin A*, which is required for cell cycle progression, *Cyclin B* (*CycB*) and *aurora B*, which are required for both cell cycle progression and cytokinesis; in contrast the housekeeping gene *Actin 5C* was unaffected (**Figure 3B-C, Figure S3C-C’**) (Mathieu et al., 2013; Matias et al., 2015). We confirmed that CycB was reduced in the ovaries of *aramis* germline RNAi flies compared to *bam* germline RNAi flies by immunofluorescence (**Figure 3D-F**). These results suggest that *aramis* is required for the proper expression of key regulators of GSC abscission.

**Figure 3.**
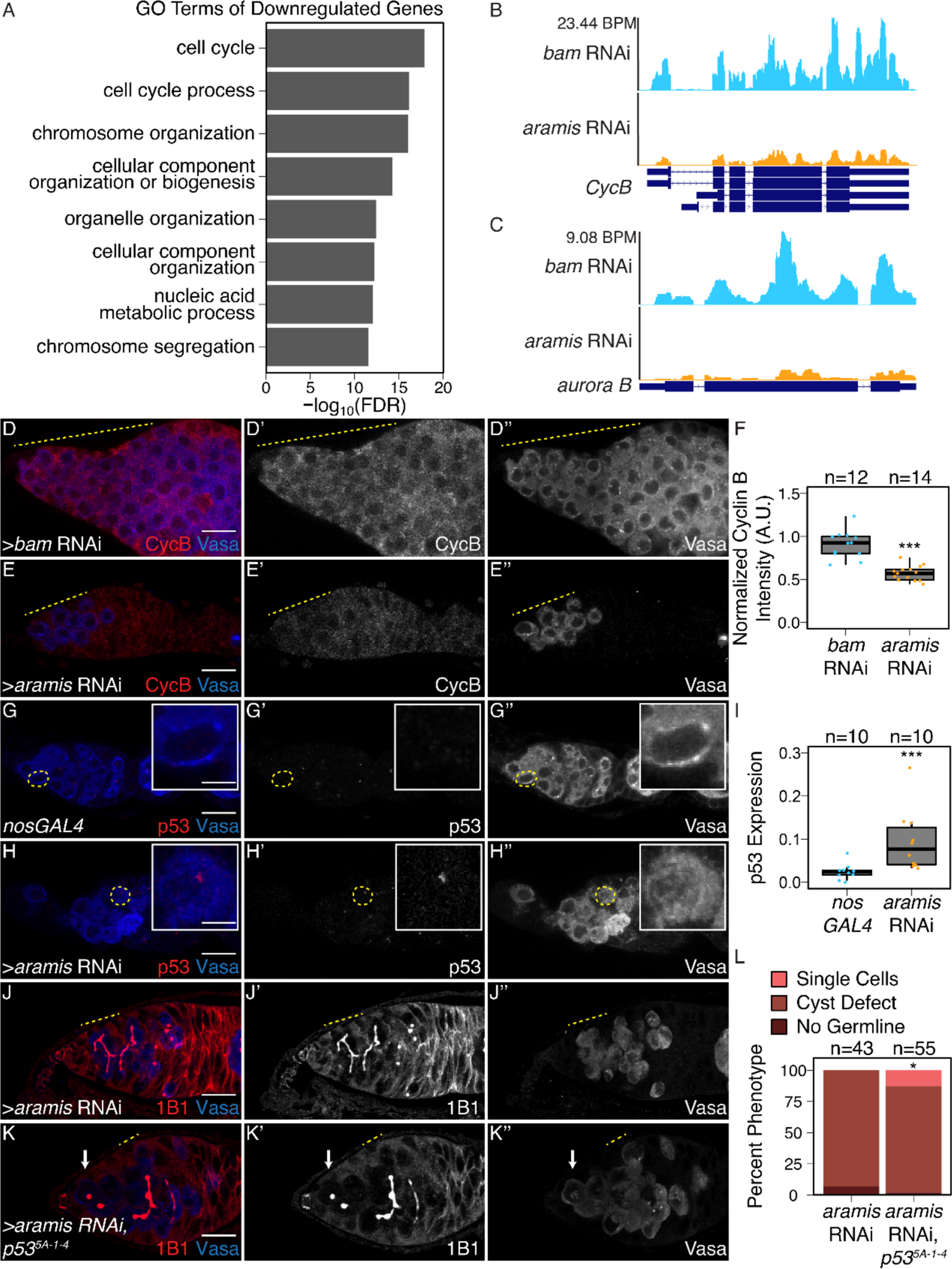
Athos, Aramis, and Porthos are required for cell cycle progression during early oogenesis. (**A**) Bar plot representing the most significant Biological Process GO terms of downregulated genes in ovaries depleted of *aramis* compared to *bam* RNAi control (FDR = False Discovery Rate from p-values using a Fisher’s exact test). (**B-C**) Genome browser tracks representing the gene locus of (**B**) *CycB* and (**C**) a*urora B* in ovaries depleted of *aramis* compared to the developmental control, *bam* RNAi. Y-axis represents the number of reads mapping to the locus in bases per million (BPM). (**D-E’’**) Confocal images of germaria stained for CycB (red, left grayscale) and Vasa (blue, right grayscale) in (**D-D’’’**) *bam* RNAi control ovaries and (**E-E’’**) *aramis* germline RNAi. (**F**) Boxplot of CycB intensity in the germline normalized to Cyc B intensity in the soma in *bam* RNAi and *aramis* RNAi (n=12-14 germaria per sample, *** = p< 0.001, Welch t-test. (**G-H’’**) Confocal images of germaria stained for p53 (red, left grayscale) and Vasa (blue, right grayscale) in (**G-G’’**) *nosGAL4*, driver control ovaries and (**H-H’’**) germline depletion of *aramis*. Cells highlighted by a dashed yellow circle represent cell shown in the inset. Driver control *nosGAL4* ovaries exhibit attenuated p53 expression in GSCs and CBs, but higher expression in cyst stages as previously reported, while p53 punctate are visible in the germline of *aramis* RNAi in the undifferentiated cells. (**I**) Box plot of percentage of pixel area exceeding the background threshold for p53 in GSCs and CBs in driver control *nosGAL4* ovaries and the germline of *aramis* RNAi indicates p53 expression is elevated in the germline over the GSCs/CBs of control ovaries. (n=10 germaria per sample, *** = p < 0.001, Welch’s t-test. (**J-K’’**) Confocal images of germaria stained for 1B1 (red, left grayscale) and Vasa (blue, right grayscale) in (**J-J’’**) germline *aramis* RNAi in a wild type background and (**K-K’’**) germline *aramis* RNAi with a mutant, null, *p53^5-A-14^* background showing presence of spectrosomes upon loss of p53. (**L**) Quantification of stem cyst phenotypes demonstrates a significant rescue upon of loss of *p53^5-A-14^* in *aramis* germline depletion compared to the wild type control (n=43-55 germaria per genotype, df=2, Fisher’s exact test p< 0.05). Scale bar for main images is 15 micron, scale bar for insets is 3.75 micron.

CycB is expressed during G2 phase after asymmetric cell division to promote GSC abscission (Flora et al., 2018a; Mathieu et al., 2013). To test if the loss of germline *aramis* leads to GSC abscission defects due to diminished expression of CycB, we attempted to express a functional CycB::GFP fusion protein in the germline under the control of a UAS/GAL4 system (**Figure S3D-D’**) (Mathieu et al., 2013). Unexpectedly, the CycB::GFP fusion protein was not expressed in the *aramis-*depleted germline, unlike the wild type (WT) germline (**Figure S3E-E’**) (Glotzer et al., 1991; Mathieu et al., 2013; Zielke et al., 2014). We considered the possibility that progression into G2 is blocked in the absence of *aramis*, precluding expression of CycB. To monitor the cell cycle, we used the Fluorescence Ubiquitin-based Cell Cycle Indicator (FUCCI) system. *Drosophila* FUCCI utilizes a GFP-tagged degron from E2f1 to mark G2, M, and G1 phases and an RFP-tagged degron from CycB to mark S, G2, and M phases (Zielke et al., 2014). We observed cells in different cell cycle stages in both WT and *bam*-depleted germaria, but the *aramis*-depleted germaria expressed neither GFP nor RFP (**Figure S3F-H’’**). Double negative reporter expression is thought to indicate early S phase, when expression of E2f1 is low and CycB is not expressed (Hinnant et al., 2017). The inability to express FPs is not due to a defect in translation as *aramis*-depleted germline can express GFP that is not tagged with the degron (**Figure S3I-I’**). Taken together, we infer that loss of *aramis* blocks cell cycle progression around late G1 phase/early S phase and prevents progression to G2 phase, when GSCs abscise from CBs.

In mammals, cells defective for ribosome biogenesis stabilize p53, which is known to impede the G1 to S transition (Agarwal et al., 1995; Senturk and Manfredi, 2013). Therefore, we hypothesized that the reduced ribosome biogenesis in the *aramis*-depleted germline leads to p53 stabilization in undifferentiated cells, driving cell cycle arrest and GSC abscission defects. To test this hypothesis, we detected p53 and Vasa in the germline by immunostaining. A hybrid dysgenic cross that expresses p53 in undifferentiated cells was utilized as a positive control, and *p53* null flies were used as negative controls (**Figure S3J-K**) (Moon et al., 2018). In WT, we observed p53 expression in the meiotic stages of germline but p53 expression in GSCs and CBs was attenuated as previously reported (**Figure 3G-G’’**) (Lu et al., 2010). However, compared to WT GSCs/CBs, we observed p53 expression in the stem cysts of the *aramis-*depleted germline (**Figure 3G-I**). Similarly, we observed p53 expression in the stem cysts of *athos-* and *porthos-*depleted germlines **(Figure S3L-M)**, further supporting that reduced ribosome biogenesis stabilizes p53. To determine if p53 stabilization is required for the cell cycle arrest in *aramis-*depleted germline cysts, we depleted *aramis* in the germline of *p53* mutants. We observed a partial but significant alleviation of the cyst phenotype, such that spectrosomes were restored (**Figure 3J-L**). This finding indicates that p53 contributes to cytokinesis failure upon loss of *aramis*, but that additional factors are also involved. Taken together, we find that *aramis-*depleted germ cells display reduced ribosome biogenesis, aberrant expression of p53 protein and a block in cell cycle progression. Reducing p53 partially alleviates the cell cycle block and GSC cytokinesis defect.

### Aramis promotes translation of Non1, a negative regulator of p53, linking ribosome biogenesis to the cell cycle

Although p53 protein levels were elevated upon loss of *aramis* in the germline, *p53* mRNA levels were not significantly altered (log_2_ fold change: −0.49; FDR: 0.49). Given that ribosome biogenesis is affected, we considered that translation of p53 or one of its regulators was altered in *aramis*-depleted germlines. To test this hypothesis, we performed polysome-seq of gonads enriched for GSCs or CBs as developmental controls, as well as gonads depleted for *aramis* in the germline (Flora et al., 2018b). We plotted the ratios of polysome-associated RNAs to total RNAs (**Figure 4A-A’’, Supplemental Table 3**) and identified 87 mRNAs with a reduced ratio upon depletion of *aramis*, suggesting that they were translated less efficiently compared to developmental controls. Loss of *aramis* reduced the levels of these 87 downregulated transcripts in polysomes, without significantly affecting their total mRNA levels (**Figure 4B, Figure S4A-A’**). These 87 transcripts encode proteins mostly associated with translation including ribosomal proteins (**Figure 4C**). To validate that Aramis regulates translation of these target mRNAs, we utilized a reporter line for the Aramis-regulated transcript encoding Ribosomal protein S2 (RpS2) that is expressed in the context of the endogenous promoter and regulatory sequences (Buszczak et al., 2007; Zhang et al., 2014). We observed reduced levels of RpS2::GFP in germlines depleted of *aramis* but not in those depleted of *bam* (**Figure 4D-F**). To ensure that reduced RpS2::GFP levels did not reflect a global decrease in translation, we visualized nascent translation using O-propargyl-puromycin (OPP). OPP is incorporated into nascent polypeptides and can be detected using click-chemistry (Sanchez et al., 2016). We observed that global translation in the germlines of ovaries depleted of *aramis* was not reduced compared to *bam* (**Figure 4G-I**). Thus, loss of *aramis* results in reduced translation of a subset of transcripts.

**Figure 4.**
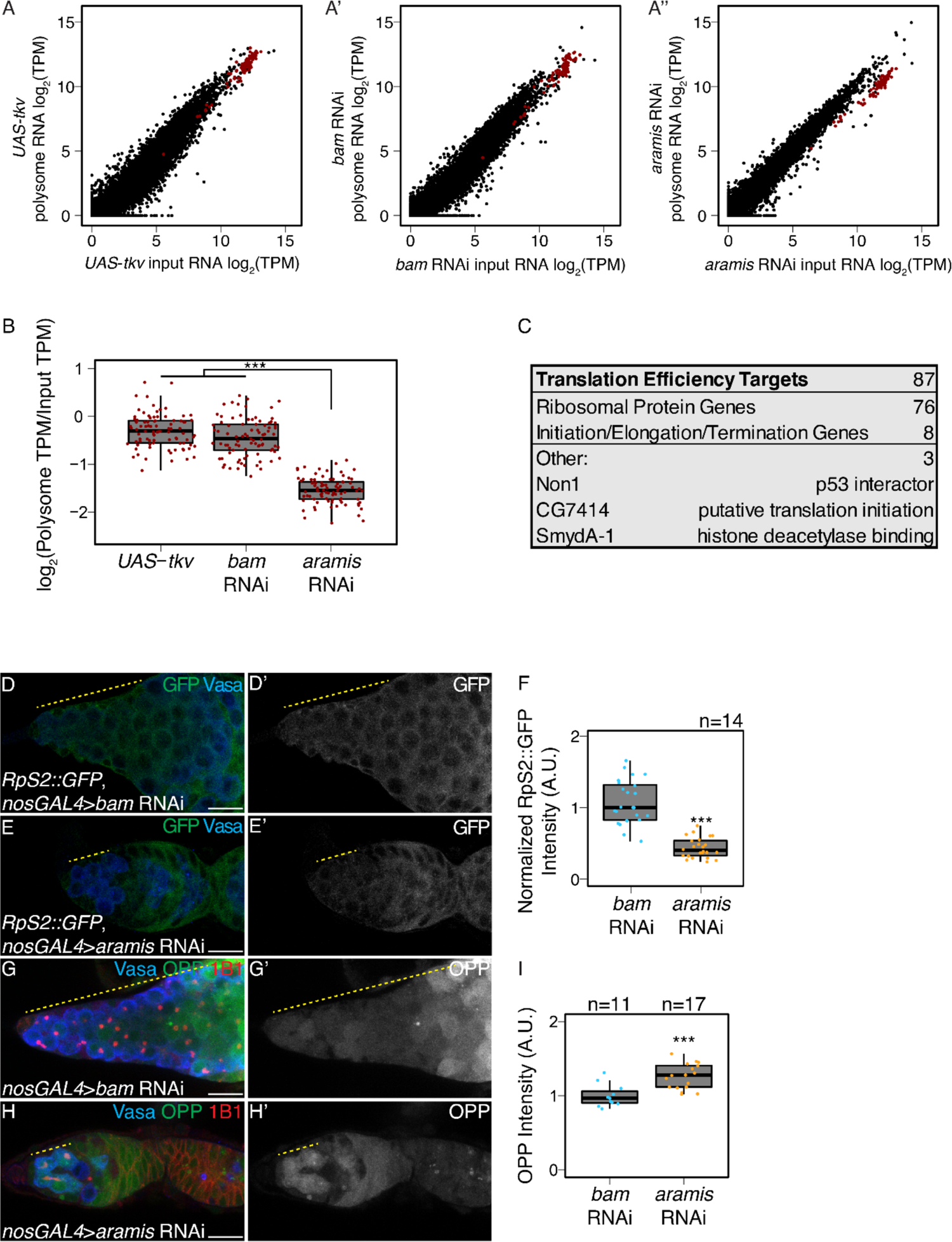
Aramis is required for efficient translation of a subset of mRNAs. (**A-A’’**) Biplots of poly(A)+ mRNA Input versus polysome associated mRNA from (**A**) ovaries genetically enriched for GSCs (*UAS-tkv*), (**A’**) Undifferentiated GSC daughter cells (*bam* RNAi) or (**A’’**) germline *aramis* RNAi ovaries. (**B**) Boxplot of translation efficiency of target genes in *UAS*-*tkv*, *bam* RNAi, and *aramis* RNAi samples (ANOVA p<0.001, post-hoc Welch’s t-test, n=87, *** = p < 0.001). (**C**) Summary of downregulated target genes identified from polysome-seq. (**D-E’**) Confocal images of germaria stained for 1B1 (red), RpS2::GFP (green, grayscale), and Vasa (blue) in (**D-D’**) *bam* RNAi control and (**E-E’**) *aramis* RNAi (yellow dashed line marks approximate region of germline used for quantification). (**F**) A.U. quantification of germline RpS2::GFP expression normalized to RpS2::GFP expression in the surrounding soma in undifferentiated daughter cells of *bam* RNAi compared to *aramis* RNAi. RpS2::GFP expression is significantly lower in *aramis* RNAi compared to control (n=14 germaria per sample, Welch’s t-test, *** = p < 0.001). (**G-H’**) Confocal images of germaria stained for 1B1 (red), OPP (green, grayscale), and Vasa (blue) in (**G-G’**) *bam* RNAi and (**H-H’**) *aramis* RNAi (yellow dashed line marks approximate region of germline used for quantification). (**I**) A.U. quantification of OPP intensity in undifferentiated daughter cells in *bam* RNAi and *aramis* RNAi (n=11-17 germaria per genotype, Welch’s t-test, *** = p < 0.001). OPP intensity is not downregulated in *aramis* RNAi compared to the control. Scale bar for all images is 15 micron.

None of these 87 translational targets have been implicated in directly controlling abscission (Mathieu et al., 2013; Matias et al., 2015). However, we noticed that the mRNA encoding Novel Nucleolar protein 1 (Non1/CG8801) was reduced in polysomes upon loss of *aramis* in the germline (**Figure 4C**). The human ortholog of Non1 is GTP Binding Protein 4 (GTPBP4), and these proteins are known to physically interact with p53 in both *Drosophila* and human cells and have been implicated in repressing p53 (mentioned as CG8801 in Lunardi et al.) (Li et al., 2018; Lunardi et al., 2010). To determine if translation of Non1 is reduced upon depletion of *aramis,* we monitored the abundance of Non1::GFP, a transgene that is under endogenous control (Sarov et al., 2016), and found that Non1::GFP was expressed in the undifferentiated GSCs and CBs (**Figure 5A-A’’**). Non1::GFP levels were reduced in the *aramis-*depleted stem cysts compared to the CBs that accumulated upon *bam*-depletion (**Figure 5B-D**), suggesting that Aramis and ribosome biogenesis promote efficient translation of Non1.

**Figure 5.**
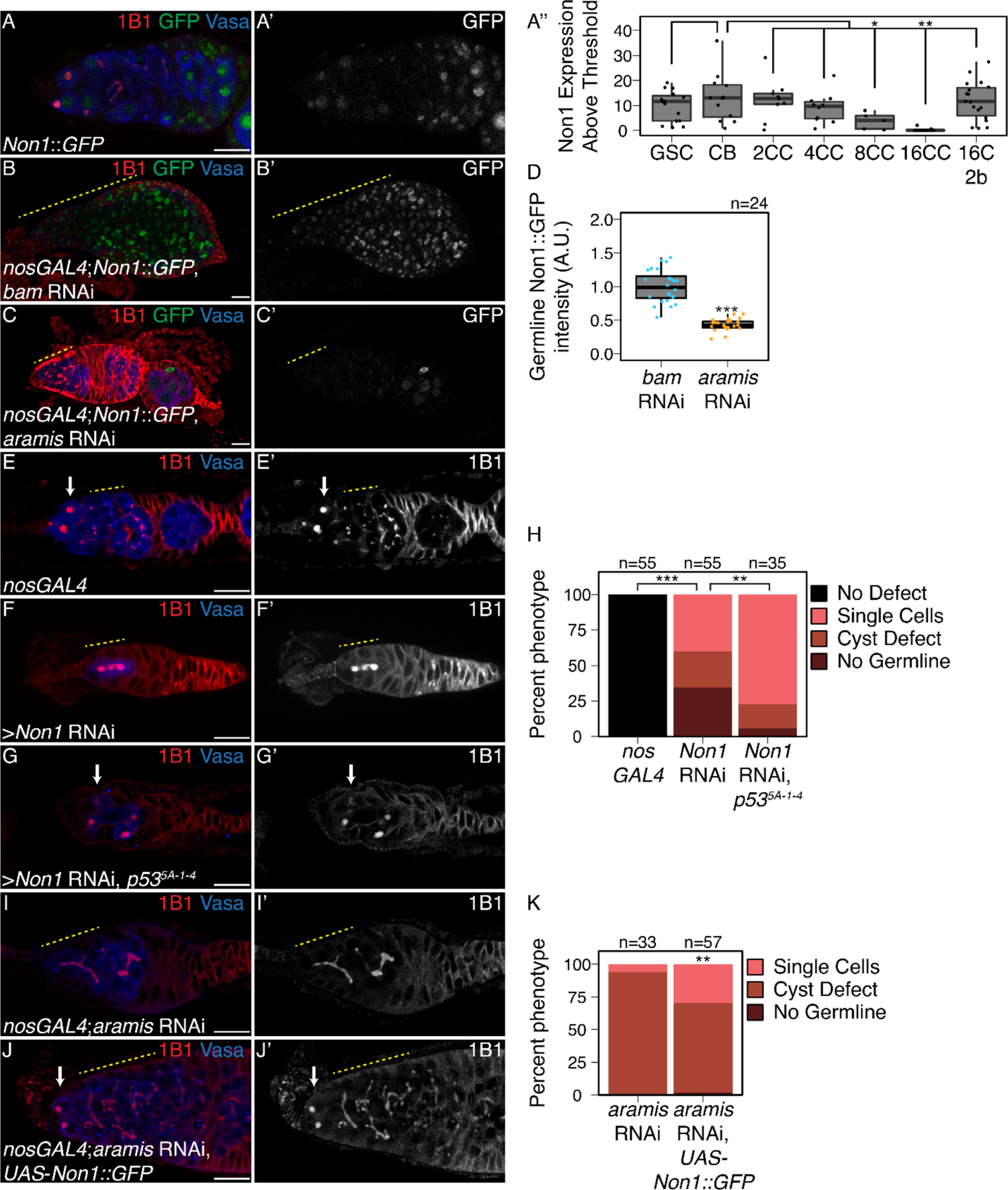
Non1 represses p53 expression to allow for GSC differentiation. (**A-A’**) Confocal images of Non1::GFP germaria stained for 1B1 (red), GFP (green, grayscale), and Vasa (blue). (**A’’**) Boxplot of Non1::GFP expression over germline development in GSCs, CBs and Cyst (CC) stages (n=5-25 cysts of each type, * = p < 0.05, ** = p < 0.01, ANOVA with Welch’s post-hoc tests). (**B-C’**) Confocal images of (**B-B’**) *bam* RNAi and (**C-C’**) *aramis* RNAi germaria both carrying *Non1::GFP* transgene stained for 1B1 (red), Vasa (blue), and Non1::GFP (green, grayscale). Yellow dashed line marks region of germline used for quantification. (**D**) Boxplot of Non1::GFP expression in the germline normalized to somatic Non1::GFP expression in *bam* RNAi and *aramis* RNAi (n=24 germaria per genotype, Welch’s t-test, *** = p < 0.001). Non1 expression is significantly lower in the germline of *aramis* RNAi compared to *bam* RNAi control. (**E-G’**) Confocal images of germaria stained for 1B1 (red, grayscale), and Vasa (blue) in (**E-E’**) *nosGAL4*, driver control ovaries, (**F-F’**) germline *Non1* RNAi, and (**G-G’**) germline *Non1* RNAi in a *p53^5-A-1-4^* background. Arrow marks the presence of a single cell (**E, G**), yellow dashed line marks a stem cyst emanating from the niche (**F-F’**) or the presence of proper cysts (**E-E’**). (**H**) Quantification of percentage of germaria with no defect (black), presence of single cell (salmon), presence of a stem cyst emanating from the niche (brown-red), or germline loss (dark red) demonstrates a significant rescue of stem cyst formation upon of loss of *Non1* in *p53^5-A-14^* compared to the *p53* wild type control (n=35-55 germaria per genotype, df=3, Fisher’s exact test with Holm-Bonferroni correction ** = p< 0.01, *** = p< 0.001). (**I-J’**) Confocal images of germaria stained for 1B1 (red, grayscale), and Vasa (blue) in (**I-I’**) *aramis* germline RNAi exhibiting stem cyst phenotype (yellow dashed line) and (**J-J**’) *aramis* germline RNAi with *Non1* overexpression exhibiting single cells (arrow). (**K**) Phenotypic quantification of *aramis* RNAi with *Non1* overexpression demonstrates a significant alleviation of the stem cyst phenotype (n=33-57 germaria per genotype, df=2, Fisher’s exact test, ** = p< 0.01). Scale bar for all images is 15 micron.

During normal oogenesis, p53 protein is expressed in cyst stages in response to recombination-induced double strand breaks (Lu et al., 2010). We found that Non1 was highly expressed at undifferentiated stages and in two- and four-cell cysts when p53 protein levels were low, whereas its expression was attenuated at eight- and 16-cell cyst stages when p53 protein levels were high (**Figure 5A-A’’, Figure S5A-B’**). Non1 was highly expressed in egg chambers, which express low levels of p53 protein suggesting that Non1 could regulate p53 protein levels. To determine if Non1 regulates GSC differentiation and p53, we depleted *Non1* in the germline. We found that germline-depletion of *Non1* results in stem cyst formation and loss of later stages, as well as increased p53 expression, phenocopying germline-depletion of *aramis*, *athos*, and *porthos* (**Figure 5E-F, H, Figure S5C-E**). In addition, we found that loss of *p53* from *Non1-*depleted germaria partially suppressed the phenotype (**Figure 5F-H**). Thus, *Non1* is regulated by *aramis* and is required for p53 suppression, cell cycle progression, and GSC abscission.

To determine if Aramis promotes GSC differentiation via translation of Non1, we restored *Non1* expression in germ cells depleted of *aramis*. Briefly, we cloned *Non1* with heterologous UTR elements under the control of the UAS/GAL4 system (see Methods) (Rørth, 1998). We found that restoring *Non1* expression in the *aramis-*depleted germline significantly attenuated the stem cysts and increased the number of cells with spectrosomes (**Figure 5I-K**). Taken together, we conclude that Non1 can partially suppress the cytokinesis defect caused by germline *aramis* depletion.

### Aramis-regulated targets contain a TOP motif in their 5’UTR

We next asked how *aramis* and efficient ribosome biogenesis promote the translation of a subset of mRNAs, including *Non1*, to regulate GSC differentiation. We hypothesized that the 87 mRNA targets share a property that make them sensitive to rRNA and ribosome levels. To identify shared characteristics, we performed *de novo* motif discovery of target genes compared to non-target genes (Heinz et al., 2010) and identified a polypyrimidine motif in the 5’UTRs of most target genes (UCUUU; E-value: 6.6e^-094^). This motif resembles the previously described TOP motif at the 5’ end of mammalian transcripts (Philippe et al., 2018; Thoreen et al., 2012). Although the existence of TOP-containing mRNAs in *Drosophila* has been speculated, to our knowledge their presence has not been explicitly demonstrated (Chen and Steensel, 2017; Qin et al., 2007). This observation motivated us to precisely determine the 5’ end of transcripts, so we analyzed previously published <u>c</u>ap <u>a</u>nalysis of <u>g</u>ene <u>e</u>xpression sequencing (CAGE-seq) data that had determined transcription start sites (TSS) in total mRNA from the ovary (**Figure 6A, Figure S6A-A’**) (Boley et al., 2014; Chen et al., 2014; dos Santos et al., 2015). Of the 87 target genes, 76 had sufficient expression in the CAGE-seq dataset to define their TSS. We performed motif discovery using the CAGE-seq data and found that 72 of 76 Aramis-regulated mRNAs have a polypyrimidine motif that starts within the first 50 nt of their TSS (**Figure 6B-C**). In mammals, it was previously thought that the canonical TOP motif begins with an invariant ‘C’ (Meyuhas, 2000; Philippe et al., 2020). However, systematic characterization of the sequence required in order for an mRNA to be regulated as a TOP containing mRNA revealed that TOP mRNAs can start with either a ‘C’ or a ‘U’ (Philippe et al., 2020). Thus, mRNAs whose efficient translation is dependent on *aramis* share a terminal polypyrimidine-rich motif in their 5’UTR that resembles a TOP motif.

**Figure 6.**
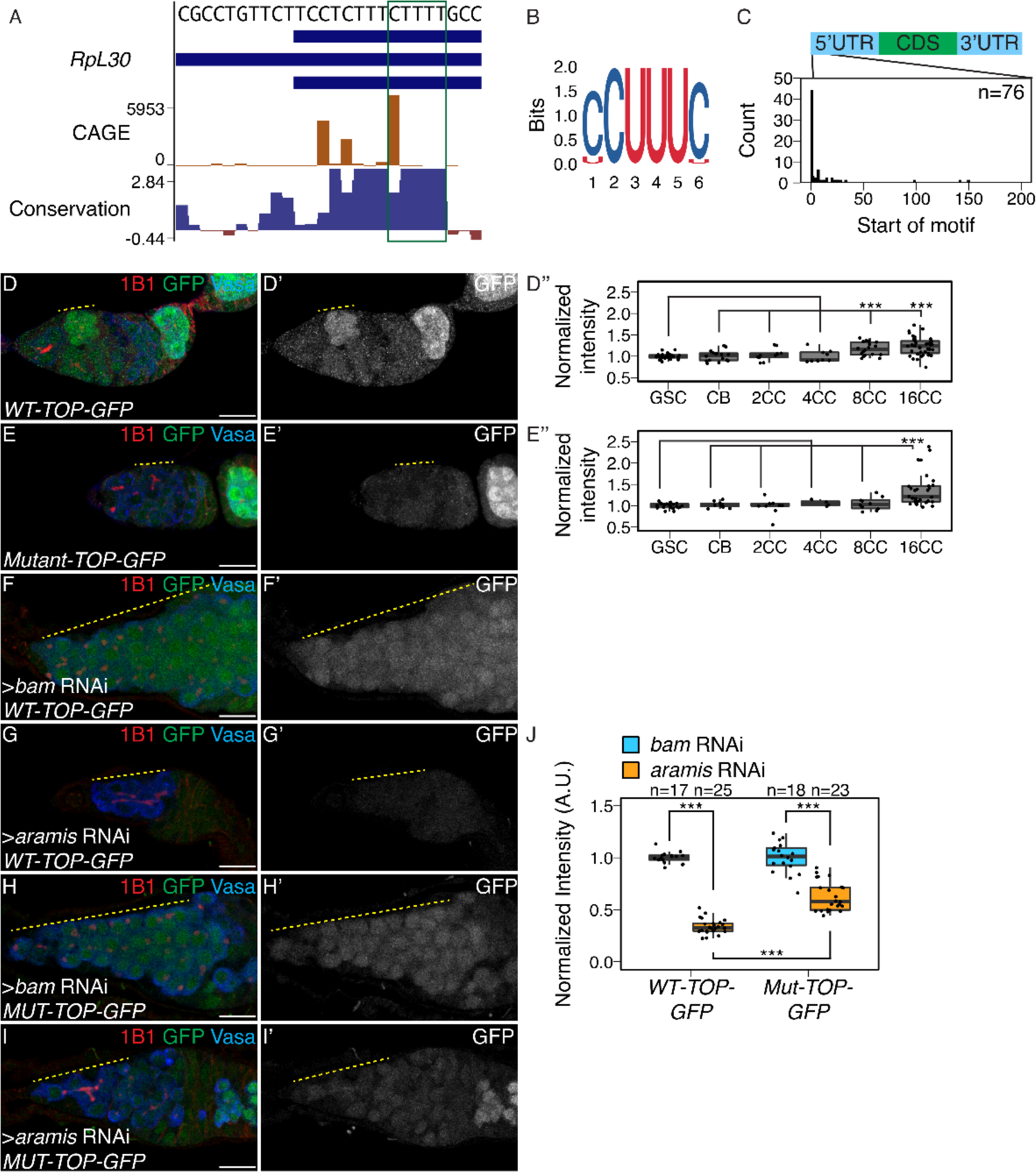
Aramis regulated mRNAs contain a TOP motif. (**A**) Genome browser tract of *RpL30* locus in ovary CAGE-seq data showing the proportion of transcripts that are produced from a given TSS (orange). Predominant TSSs are shown in orange and putative TOP motif indicated with a green box. The bottom blue and red graph represents sequence conservation of the locus across *Diptera*. The dominant TSS initiates with a canonical TOP motif. (**B**) Sequence logo generated from *de novo* motif discovery on the first 200 bases downstream of CAGE derived TSSs of *aramis* translation target genes resembles a canonical TOP motif. (**C**) Histogram representing the location of the first 5-mer polypyrimidine sequence from each CAGE based TSS of *aramis* translation target genes demonstrates that the TOP motifs occur proximal to the TSS (n=76 targets). (**D-E’’**) Confocal images and quantifications of (**D-D’**) *WT-TOP-GFP* and (**E-E’**) *Mut-TOP-GFP* reporter expression stained for 1B1 (red), GFP (green, grayscale), and Vasa (blue). Yellow dotted-line marks increased reporter expression in 8-cell cysts of *WT-TOP-GFP* but not in *Mut-TOP-GFP*. Reporter expression was quantified over germline development for (**D’’**) *WT-TOP-GFP* and (**E’’**) *Mut-TOP-GFP* reporter expression and normalized to expression in the GSC reveals dynamic expression based on the presence of a TOP motif. (**F-G’**) Confocal images of *WT-TOP-GFP* reporter ovarioles showing 1B1 (red), GFP (green, grayscale), and Vasa (blue) in (**F-F’**) *bam* germline depletion as a developmental control and (**G-G’**) *aramis* germline depleted ovaries. Yellow dotted lines indicate germline. (**H-I’**) Confocal images of *Mut-TOP-GFP* reporter expression showing 1B1 (red), GFP (green, grayscale), and Vasa (blue) in (**H-H’**) *bam* RNAi and (**I-I’**) *aramis* germline RNAi. Yellow dotted lines indicate germline. (**J**) A.U. quantification of WT and Mutant TOP reporter expression in undifferentiated daughter cells in *bam* RNAi compared *aramis* RNAi demonstrates that the *WT-TOP-GFP* reporter shows significantly lower expression in *aramis* RNAi than the *Mut-TOP-GFP* relative to the expression of the respective reporters in *bam* RNAi (n=17-25 germaria per genotype, with Welch’s t-test *** = p<0.001). Scale bar for all images is 15 micron.

In vertebrates, most canonical TOP-regulated mRNAs encode ribosomal proteins and translation initiation factors that are coordinately upregulated in response to growth cues mediated by the Target of Rapamycin (TOR) pathway and the TOR complex 1 (mTORC1) (Hornstein et al., 2001; Iadevaia et al., 2014; Kim et al., 2008; Meyuhas and Kahan, 2015; Pallares-Cartes et al., 2012) Indeed, 76 of the 87 Aramis targets were ribosomal proteins, and 9 were known or putative translation factors, consistent with TOP-containing RNAs in vertebrates (**Figure 4C**, **Supplemental Table 4**). To determine if the putative TOP motifs that we identified are sensitive to TORC1 activity, we designed “TOP reporter” constructs. Specifically, the germline-specific *nanos* promoter was employed to drive expression of an mRNA with 1) the 5’UTR of the *aramis* target *RpL30*, which contains a putative TOP motif, 2) the coding sequence for a GFP-HA fusion protein and 3) a 3’UTR (K10) that is not translationally repressed (Flora et al., 2018b; Serano et al., 1994), referred to as the WT-TOP reporter (**Figure S6B**). As a control, we created a construct in which the polypyrimidine sequence was mutated to a polypurine sequence referred to as the Mut-TOP reporter (**Figure S6B**).

In *Drosophila*, TORC1 activity increases in 8- and 16-cell cysts (Hong et al., 2012; Kim et al., 2017). We found that the WT-TOP reporter displayed peak expression in 8-cell cysts, whereas the Mutant-TOP reporter did not (**Figure 6D-E’’**), suggesting that the WT-TOP reporter is sensitive to TORC1 activity. Moreover, depletion of *Nitrogen permease regulator-like 3* (*Nprl3*), an inhibitor of TORC1 (Wei et al., 2014), led to a significant increase in expression of the WT-TOP reporter but not the Mutant-TOP reporter **(Figure S6C-G)**. Additionally, to attenuate TORC1 activity, we depleted *raptor*, one of the subunits of the TORC1 complex (Hong et al., 2012; Loewith and Hall, 2011). Here we found that the WT-TOP reporter had a significant decrease in reporter expression while the Mutant-TOP reporter did not show a decrease in expression **(Figure S6H-L)**. Taken together, our data suggest that Aramis-target transcripts contain TOP motifs that are sensitive to TORC1 activity. However, we note that our TOP reporter did not recapitulate the pattern of Non1::GFP expression, suggesting that Non1 may have additional regulators that modulate its protein levels in the cyst stages.

TOP mRNAs show increased translation in response to TOR signaling, leading to increased ribosome biogenesis (Jefferies et al., 1997; Jia et al., 2021; Powers and Walter, 1999; Thoreen et al., 2012). However, to our knowledge, whether reduced ribosome biogenesis can coordinately diminish the translation of TOP mRNAs to balance and lower ribosome protein production and thus balance the levels of the distinct components needed for full ribosome assembly is not known. To address this question, we crossed the transgenic flies carrying the WT-TOP reporter and Mutant-TOP reporter into *bam* and *aramis* germline RNAi backgrounds. We found that the expression from the WT-TOP reporter was reduced by 2.9-fold in the germline of *aramis* RNAi ovaries compared to *bam* RNAi ovaries (**Figure 6F-G, J**). In contrast, the Mutant-TOP reporter was only reduced by 1.6-fold in the germline of *aramis* RNAi ovaries compared to *bam* RNAi ovaries (**Figure 6H-J**). This suggests that the TOP motif-containing mRNAs are sensitive to ribosome biogenesis.

### Larp binds TOP sequences in *Drosophila*

Next, we sought to determine how TOP-containing mRNAs are regulated downstream of Aramis. In mammalian cells, Larp1 is a critical negative regulator of TOP-containing RNAs during nutrient deprivation (Berman et al., 2020; Fonseca et al., 2015; Hong et al., 2017; Philippe et al., 2020; Tcherkezian et al., 2014). Therefore, we hypothesized that *Drosophila* Larp reduces the translation of TOP-containing mRNAs when rRNA biogenesis is reduced upon loss of *aramis*. First, using an available gene-trap line in which endogenous Larp is tagged with GFP and 3xFLAG, we confirmed that Larp was robustly expressed throughout all stages of oogenesis including in GSCs **(Figure S7A-A’)**.

Next, we performed electrophoretic mobility shift assays (EMSA) to examine protein-RNA interactions with purified *Drosophila* Larp-DM15, the conserved domain that binds to TOP sequences in vertebrates (Lahr et al., 2017). As probes, we utilized capped 42-nt RNAs corresponding to the 5’UTRs of *RpL30* and *Non1*, including their respective TOP sequences. We observed a gel shift with these RNA oligos in the presence of increasing concentrations of Larp-DM15 (**Figure 7A-A’, Figure S7B**), and this shift was abrogated when the TOP sequences were mutated to purines **(Figure S7C-C’).** To determine if Larp interacts with TOP-containing mRNAs *in vivo*, we immunopurified Larp::GFP::3xFLAG from the ovaries of the gene-trap line and performed RNA-seq (**Figure S7D**). We uncovered 156 mRNAs that were bound to Larp, and 84 of these were among the 87 *aramis* translationally regulated targets, including *Non1*, *RpL30*, and *RpS2* (**Figure 7B-C**, **Supplemental Table 5**). Thus, *Drosophila* Larp binds to TOP sequences *in vitro* and TOP-containing mRNAs *in vivo*.

**Figure 7.**
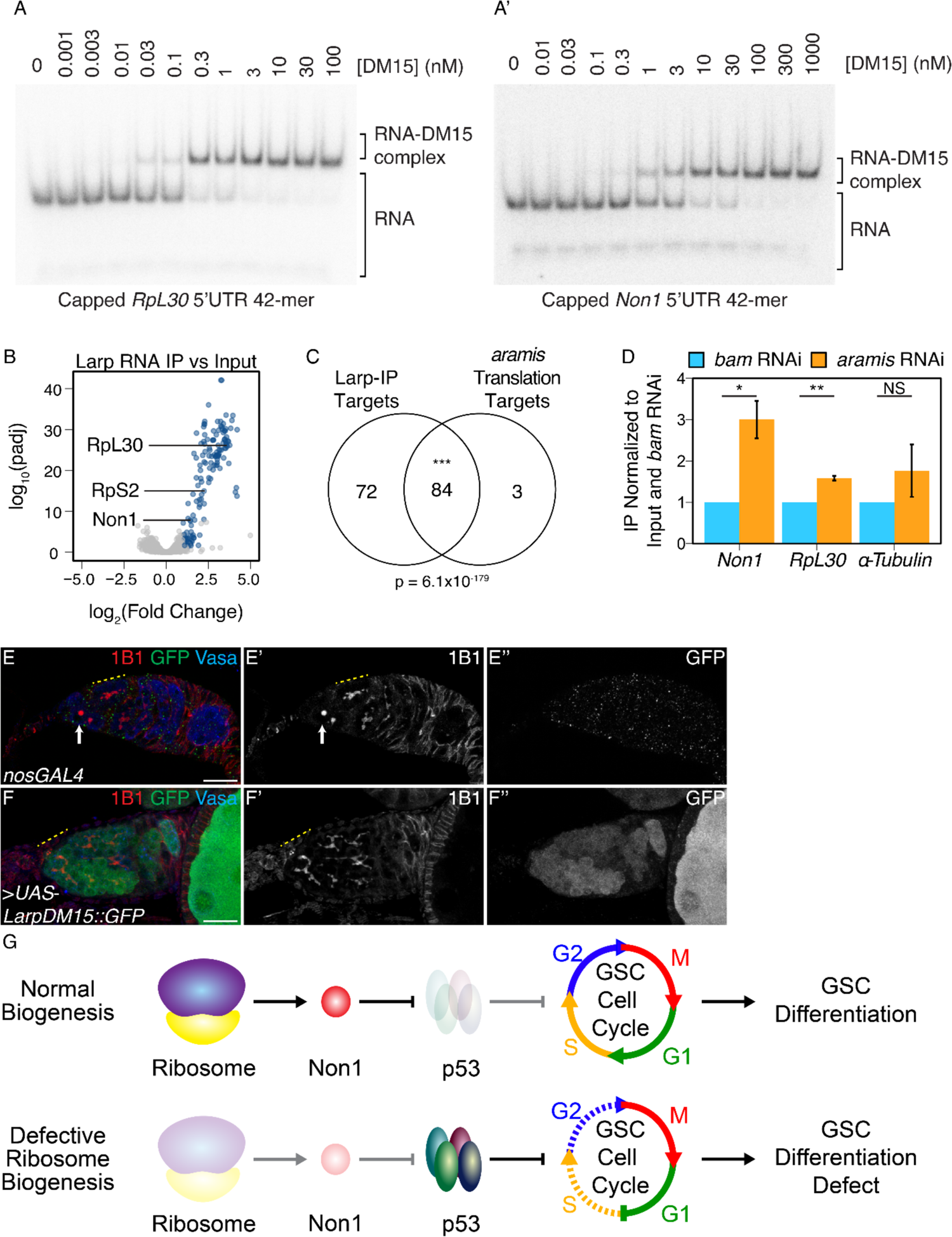
Larp binds to TOP mRNAs and binding is regulated by Aramis. (**A-A’**) EMSA of Larp-DM15 and the leading 42 nucleotides of (**A**) *RpL30* and (**A’**) *Non1* with increasing concentrations of Larp-DM15 from left to right indicates that both RNAs bind to Larp-DM15. (**B**) Volcano plot of mRNAs in Larp::GFP::3xFLAG IP compared to input. Blue points represent mRNAs significantly enriched in Larp::GFP::3xFLAG compared to input, but not enriched in an IgG control compared to input. (**C**) Venn diagram of overlapping Larp IP targets and *aramis* RNAi polysome seq targets indicates that Larp physically associates with mRNAs that are translationally downregulated in germline *aramis* RNAi (p < 0.001, Hypergeometric Test). (**D**) Bar plot representing the fold enrichment of mRNAs from Larp RNA IP in germline *aramis* RNAi relative to matched *bam* RNAi ovaries as a developmental control measured with qPCR (n=3, * = p<0.5, ** = p<0.01, NS = nonsignificant, One-sample t-test, mu=1) indicates that more of two *aramis* translation targets *Non1* and *RpL30* are bound by Larp in *aramis* RNAi. (**E-F’’**) Confocal images of (**E-E’’**) *nosGAL4*, driver control and (**F-F’’**) ovaries overexpressing the DM15 region of Larp in the germline ovaries stained for 1B1 (red, left grayscale), Vasa (blue), and Larp-DM15::GFP (green, right grayscale). Overexpression of Larp results in an accumulation of extended 1B1 structures (highlighted with a dotted yellow line), marking interconnected cells when Larp-DM15 is overexpressed compared to *nosGAL4*, driver control ovaries. (**G**) In conditions with normal ribosome biogenesis Non1 is efficiently translated, downregulating p53 levels allowing for progression through the cell cycle. When ribosome biogenesis is perturbed Non1 is not translated to sufficient levels, resulting in the accumulation of p53 and cell cycle arrest. Scale bar for all images is 15 micron.

To test our hypothesis that *Drosophila* Larp inhibits the translation of TOP-containing mRNAs upon loss of *aramis*, we immunopurified Larp::GFP::3xFLAG from germline *bam* RNAi ovaries and germline *aramis* RNAi ovaries. Larp protein is not expressed at higher levels in *aramis* RNAi compared to developmental control *bam* RNAi (**Figure S7E-G**). We found that Larp binding to *aramis* target mRNAs *Non1* and *RpL30* was increased in *aramis* RNAi ovaries compared to *bam* RNAi ovaries (**Figure 7D, Figure S7H**). In contrast, a non-target mRNA that does not contain a TOP motif, *alpha-tubulin* mRNA, did not have a significant increase in binding to Larp in *aramis* RNAi ovaries compared to *bam* RNAi ovaries. Overall, these data suggest that reduced rRNA biogenesis upon loss of *aramis* increases Larp binding to the TOP-containing mRNAs *Non1* and *RpL30*.

If loss of *aramis* inhibits the translation of TOP-containing mRNAs due to increased Larp binding, then overexpression of Larp would be expected to phenocopy germline depletion of *aramis*. Unphosphorylated Larp binds to TOP motifs more efficiently, but the precise phosphorylation sites of *Drosophila* Larp, to our knowledge, are currently unknown (Hong et al., 2017). To circumvent this issue, we overexpressed the DM15 domain of Larp which we showed binds the *RpL30* and *Non1* TOP motifs *in vitro* (**Figure 7A-A’**), and, based on homology to mammalian Larp1, lacks majority of the putative phosphorylation sites (Jia et al., 2021; Lahr et al., 2017; Philippe et al., 2018). We found that overexpression of a Larp-DM15::GFP fusion in the germline resulted in fusome-like structures extending from the niche (**Figure 7E-F’**). Additionally, ovaries overexpressing Larp-DM15 had 32-cell egg chambers, which were not observed in control ovaries (**Figure S7I-I’**). The presence of 32-cell egg chambers is emblematic of cytokinesis defects that occur during early oogenesis (Mathieu et al., 2013; Matias et al., 2015; Sanchez et al., 2016). Our findings indicate that these cells are delayed in cytokinesis and that over expression of Larp partially phenocopies depletion of *aramis*.

## Discussion

During *Drosophila* oogenesis, efficient ribosome biogenesis is required in the germline for proper GSC cytokinesis and differentiation. The outstanding questions that needed to be addressed were: 1) Why does disrupted ribosome biogenesis impair GSC abscission and differentiation? and 2) How does the GSC monitor and couple ribosome abundance to differentiation? Our results suggest that germline ribosome biogenesis defect stalls the cell cycle, resulting a loss of differentiation and the formation of stem cysts. We discovered that proper ribosome biogenesis is monitored through a translation control module that allows for co-regulation of ribosomal proteins and a p53 repressor. Loss of *aramis*, *athos* and *porthos* reduces ribosome biogenesis and inhibits translation of a p53 repressor, leading to p53 stabilization, cell cycle arrest and loss of stem cell differentiation (**Figure 7G**).

### Aramis, Athos, and Porthos are required for efficient ribosome biogenesis in *Drosophila*

We provide evidence that Aramis, Athos and Porthos play a role in ribosome biogenesis in *Drosophila*, similar to their orthologs in yeast (Bohnsack et al., 2008; Granneman et al., 2006; Khoshnevis et al., 2016; O’day et al., 1996) and mammals (Sekiguchi et al., 2006; Tafforeau et al., 2013; Zhang et al., 2011). Their role in ribosome biogenesis is likely a direct function of these helicases as they physically interact with precursor rRNA. In yeast, Rok1, the ortholog of Aramis, binds to several sites on pre-rRNA, predominantly in the 18S region (Bohnsack et al., 2008; Khoshnevis et al., 2016; Martin et al., 2014). This is consistent with the small subunit ribosome biogenesis defect we observe upon loss of *aramis* in *Drosophila*. Rrp3, the yeast ortholog of Porthos, promotes proper cleavage of pre-rRNA and is required for proper 18S rRNA production (Granneman et al., 2006; O’day et al., 1996). DDX47, the mammalian ortholog of Porthos, binds to early rRNA precursors as well as proteins involved in ribosome biogenesis (Sekiguchi et al., 2006). Consistent with these findings, we find that Aramis and Porthos promote 40S ribosome biogenesis. DHX33, the mammalian ortholog of Athos, has been implicated in facilitating rRNA synthesis (Zhang et al., 2011). In contrast, we find that Athos promotes 60S ribosome biogenesis by directly interacting with rRNA. However, we cannot rule out that Athos also affects transcription of rRNA in *Drosophila* as it does in mammals (Zhang et al., 2011). Overall, we find that each mammalian ortholog of Aramis, Athos, and Porthos has consistent ribosome subunit defects, suggesting that the function of these helicases is conserved from flies to mammals. Intriguingly, DDX52 (Aramis) is one of the 15 genes deleted in 17q12 syndrome (Hendrix et al., 2012). 17q12 syndrome results in delayed development, intellectual disability, and, more rarely, underdevelopment of organs such as the uterus (Bernardini et al., 2009; Hendrix et al., 2012). Our finding that Aramis disrupts stem cell differentiation could explain some of the poorly understood defects in 17q12 syndrome.

### Ribosome biogenesis defects leads to cell cycle defects mediated by p53

Here we report that three RNA helicases, *aramis*, *athos*, and *porthos,* that promote proper ribosome biogenesis in *Drosophila* are required in the germline for fertility. Loss of *aramis*, *athos*, and *porthos* causes formation of a “stem cyst” and loss of later stage oocytes. Stem cysts are a characteristic manifestation of ribosome biogenesis deficiency wherein GSCs are unable to complete cytokinesis and fail to express the differentiation factor Bam, which in GSCs is initiated at G2 of the cell cycle (Sanchez et al., 2016; Zhang et al., 2014). Our RNA seq and cell cycle analysis indicates that depletion of *aramis* blocks the cell cycle at G1, and that failure to progress to G2 prevents abscission and expression of Bam. Thus, our results suggest that ribosome biogenesis defects in the germline stall the cell cycle, resulting in formation of stem cysts and sterility.

In most tissues in *Drosophila,* p53 primarily activates apoptosis, however, in the germline p53 is activated during meiosis and does not cause cell death (Fan et al., 2010; Lu et al., 2010). Furthermore, p53 activation in the germline is required for germline repopulation and GSC survival after genetic insult, implicating p53 as a potential cell cycle regulator (Ma et al., 2016; Tasnim and Kelleher, 2018). Our observation that reduction of *p53* partially rescues a stem cyst defect caused by ribosome deficiency due to germline depletion of *aramis* indicates that the G1 block in GSCs is, in part, mediated by p53 activation. Thus, in *Drosophila* GSCs, p53 blocks the GSC cell cycle and is sensitive to rRNA production. The developmental upregulation of p53 during GSC differentiation concomitant with lower ribosome levels parallels observations in disease states, such as ribosomopathies (Calo et al., 2018; Deisenroth and Zhang, 2010; Pereboom et al., 2011; Yelick and Trainor, 2015).

We find that p53 levels in GSCs are regulated by conserved p53 regulator Non1. In mammalian cells, increased free RpS7 protein due to nucleolar stress binds and sequesters MDM2, a repressor of p53, freeing p53, resulting in G1 cell cycle arrest (Deisenroth and Zhang, 2010; Zhang and Lu, 2009). *Drosophila* have no identified homolog to MDM2. It is not fully known how ribosome levels are monitored in *Drosophila* in the absence of MDM2 and how this contributes to cell cycle progression. In *Drosophila,* Non1 levels are high in the GSCs and p53 is low, and reciprocally Non1 levels are low during meiosis, but p53 is expressed. Our finding that loss of Aramis leads to diminished Non1 and elevated p53, and that either loss of p53 or elevated Non1 suppress differentiation defects caused by loss of Aramis, suggests that, in the female germline, Non1 may fulfill the function of Mdm2 by promoting p53 degradation during *Drosophila* oogenesis. While Non1 has been shown to directly interact with p53, how it regulates p53 levels in both humans and *Drosophila* is not known (Li et al., 2018; Lunardi et al., 2010). Overall, our data place Non1 downstream of ribosome biogenesis and upstream of p53 in controlling cell cycle progression and GSC differentiation. However, our data do not rule out that Non1 may also act upstream of or in parallel to Aramis.

The vertebrate ortholog of Non1, GTPBP4, also controls p53 levels and is upregulated in some cancers (Li et al., 2018; Lunardi et al., 2010; Yu et al., 2016). This suggests that there may be parallel pathways for monitoring ribosome levels via p53 in different tissue types. Unlike *Drosophila* Non1, its ortholog, GTPBP4 has not been identified as a TOP mRNA, so if it similarly acts as a mediator between ribosome biogenesis and the cell cycle it is likely activated in a somewhat different manner (Philippe et al., 2020). However, mammalian Larp1 is required for proper cell cycle progression and cytokinesis (Burrows et al., 2010; Tcherkezian et al., 2014). Excitingly several differentiation and cell cycle regulation genes in mammals are TOP mRNAs regulated by Larp1, including Tumor Protein, Translationally-Controlled 1 (TPT1) and Nucleosome Assembly Protein 1 Like 1 (NAP1L1) (Philippe et al., 2020). TPT1 is a cancer associated factor that has been implicated in activating pluripotency (Burrows et al., 2010; Qiao et al., 2018). Similarly, NAP1L1, a nucleosome assembly protein, is required to maintain proper cell cycle control as loss of NAP1L1 results in cell cycle exit and premature differentiation. Overall, although the specific targets of Larp1 in mammals may differ from those in *Drosophila*, the mechanism by which Larp modulates cell cycle and differentiation may be conserved.

### Ribosome biogenesis defects leads to repression of TOP-containing mRNA

TOP-containing mRNAs are known to be coregulated to coordinate ribosome production in response to nutrition or other environmental cues (Kimball, 2002; Meyuhas and Kahan, 2015; Tang et al., 2001). Surprisingly, our observation that loss of *aramis* reduces translation of a cohort of TOP-containing mRNAs, including Non1, suggests that the TOP motif also sensitizes their translation to lowered levels of rRNA. This notion is supported by TOP reporter assays demonstrating that reduced translation upon loss of *aramis* requires the TOP motif. We hypothesize that limiting TOP mRNA translation lowers ribosomal protein production to maintain a balance with reduced rRNA production. This mechanism would prevent the production of excess ribosomal proteins that cannot be integrated into ribosomes and the ensuing harmful aggregates (Tye et al., 2019). Additionally, it would coordinate rRNA production and ribosomal protein translation during normal germline development, where it is known that the level of ribosome biogenesis and of global translation are dynamic (Blatt et al., 2020a; Fichelson et al., 2009; Sanchez et al., 2016; Zhang et al., 2014).

### Larp transduces growth status to ribosome biogenesis targets

Recent work has shown that the translation and stability of TOP-containing mRNAs are mediated by Larp1 and its phosphorylation (Berman et al., 2020; Hong et al., 2017; Jia et al., 2021). We found that perturbing rRNA production and thus ribosome biogenesis, without directly targeting ribosomal proteins, similarly results in dysregulation of TOP mRNAs. Our data show that *Drosophila* Larp binds the *RpL30* and *Non1* 5’UTR in a TOP-dependent manner *in vitro* and to nearly all of the translation targets we identified *in vivo.* Together these data suggest that rRNA production regulates TOP mRNAs via Larp. Furthermore, the cytokinesis defect caused by overexpression of Larp-DM15 in the germline suggests that Larp regulation could maintain the homeostasis of ribosome biogenesis more broadly by balancing the expression of ribosomal protein production with the rate of other aspects of ribosome biogenesis, such as rRNA processing, during development.

Previous studies indicate that unphosphorylated Larp1 binds to and represses its targets more efficiently than phosphorylated Larp1 (Fonseca et al., 2018; Hong et al., 2017; Jia et al., 2021). Thus, although we do not know the identity of the kinase that phosphorylates Larp in *Drosophila*, we hypothesize that Larp is not phosphorylated upon loss of *aramis, athos* and *porthos*, when ribosome biogenesis is perturbed. We propose that until ribosome biogenesis homeostasis is reached, this kinase will remain inactive, continuously increasing the pool of dephosphorylated Larp. In this scenario, as dephosphorylated Larp accumulates, it begins to bind its targets. Initially, it will bind its highest affinity targets, presumably encoding ribosomal proteins and repress their translation to rebalance ribosomal protein production with rRNA production. Consistent with this model, the TOP motif in *RpL30* is bound by Larp even more tightly with a nearly 9-fold higher affinity compared to the *Non1* TOP site (**Figure S7B**). We propose that such differences in affinity may allow Larp to repress ribosomal protein translation to facilitate cellular homeostasis without immediately causing cell cycle arrest. However, if homeostasis cannot be achieved and sufficient dephosphorylated Larp accumulates, Larp will also bind and repress the translation of lower affinity targets. Repression of Non1 in this manner would result in cell cycle arrest and block differentiation as occurs upon *aramis* depletion.

### Ribosome biogenesis in stem cell differentiation and ribosomopathies

Ribosomopathies arise from defects in ribosomal components or ribosome biogenesis and include a number of diseases such as Diamond-Blackfan anemia, Treacher Collins syndrome, Shwachman-Diamond syndrome, and 5q-myelodysplastic syndrome (Armistead and Triggs-Raine, 2014; Draptchinskaia et al., 1999; McGowan et al., 2011; Valdez et al., 2004; Warren, 2018). Despite the ubiquitous requirement for ribosomes and translation, ribosomopathies cause tissue-specific disease (Armistead and Triggs-Raine, 2014). The underlying mechanisms of tissue specificity remain unresolved.

In this study we demonstrate that loss of helicases involved in rRNA processing lead to perturbed ribosome biogenesis and, ultimately, cell cycle arrest. Given that *Drosophila* germ cells undergo an atypical cell cycle program as a normal part of their development it may be that this underlying cellular program in the germline leads to the tissue-specific symptom of aberrant stem cyst formation (McKearin and Spradling, 1990). This model implies that other tissues would likewise exhibit unique tissue-specific manifestations of ribosomopathies due to their underlying cell state and underscores the need to further explore tissue-specific differentiation programs and development to shed light not only on ribosomopathies but on other tissue-specific diseases associated with ubiquitous processes. Although it is also possible that phenotypic differences arise from a common molecular cause, our data suggests two sources of potential tissue specificity: 1) tissues express different cohorts of mRNAs, such as *Non1*, that are sensitive to ribosome levels. For example, we find that in *Drosophila* macrophages, RNAs that regulate the metabolic state of macrophages and influence their migration require increased levels of ribosomes for their translation (Emtenani et al., 2021). 2) p53 activation, as has been previously described, is differentially tolerated in different developing tissues (Bowen and Attardi, 2019; Calo et al., 2018; Jones et al., 2008). Together, both mechanisms could begin to explain the tissue-specific nature of ribosomopathies and their link to differentiation.

## Acknowledgements

We are grateful to all members of the Rangan and Fuchs labs for their discussion and comments on the manuscript. We also thanks Dr. Sammons, Dr. Marlow, Life Science Editors, for their thoughts and comments the manuscript Additionally, we thank the Bloomington Stock Center, the Vienna *Drosophila* Resource Center, the BDGP Gene Disruption Project, and Flybase for fly stocks, reagents, and other resources. P.R. is funded by the NIH/NIGMS (R01GM111779-06 and RO1GM135628-01), G.F. is funded by NSF MCB-2047629 and NIH RO3 AI144839, D.E.S. was funded by Marie Curie CIG 334077/IRTIM and the Austrian Science Fund (FWF) grant ASI_FWF01_P29638S, and A.B is funded by NIH R01GM116889 and American Cancer Society RSG-17-197-01-RMC.

## Author Contributions

Conceptualization, E.T.M., P.B., G.F., and P.R.; Methodology, E.T.M., P.B., G.F., and P.R.; Investigation, E.T.M., P.B., E.N., R.L., S.S., H.Y., T.P., and S.E.; Writing – Original Draft, E.T.M., D.E.S., and P.R.; Writing – Review & Editing, E.T.M., P.B., D.E.S, A.B., G.F., and P.R.; Funding Acquisition, G.F. and P.R.; Visualization, E.T.M., E.N.; Supervision, G.F. and P.R.

---

**Supplemental Figure 1.**
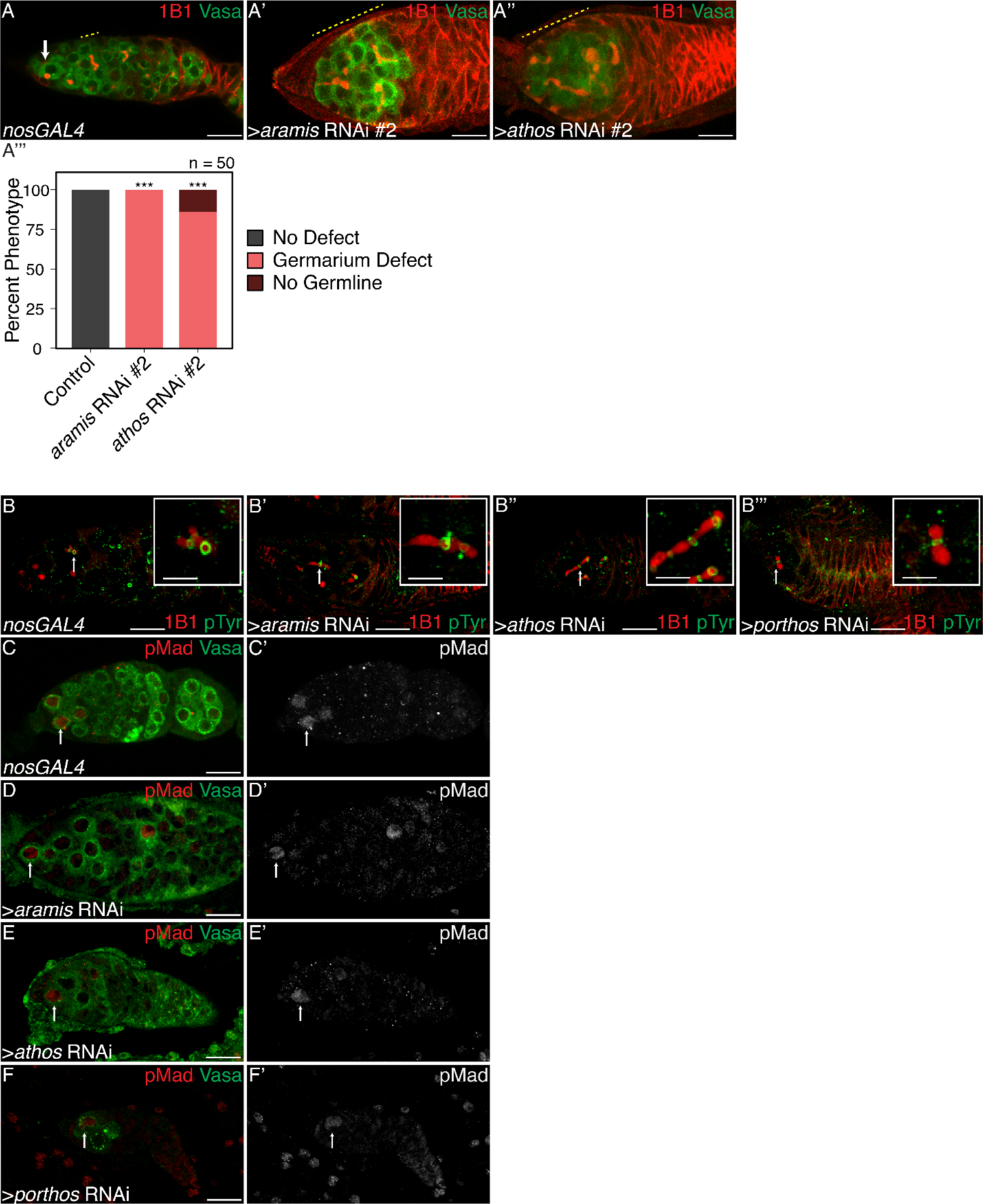
Aramis, Athos, and Porthos are required for proper cytokinesis and differentiation, related to. **Figure 1**. (**A-A’’’**) Confocal images of (**A**) *nosGAL4*, driver control and germline RNAi knockdown using additional RNAi lines for (**A’**) *aramis* and (**A’’**) *athos* stained for 1B1 (red) and Vasa (green). (**A’’’**) Quantification of percentage of germaria with no defect (black), stem cysts (salmon), or germline loss (dark red) in ovaries depleted of *athos*, *aramis*, or *porthos* compared to control ovaries recapitulates the phenotypes with independent RNAi lines (n=50, df=2, *** = p<0.001, Fisher’s exact test with Holm-Bonferroni correction). (**B-B’’’**) Confocal images of germaria stained for 1B1 (red) and Phospho-tyrosine (green). Ring canals, marked by Phosopho-tyrosine, connect differentiating cysts in (**B**) control *nosGAL4* ovaries and in between the interconnected cells of ovaries depleted of (**B’**) *athos*, (**B’’**) *aramis*, and (**B’’’**) *porthos* with 1B1 positive structures going through the ring canals. (**C-F’**) Confocal images of germaria stained for pMad (red, grayscale) and Vasa (green). In (**C**) control ovaries nuclear pMad staining occurs in cells proximal to the niche marking GSCs. Nuclear pMad staining in ovaries depleted of (**D**) *athos*, (**E**) *aramis*, and (**F**) *porthos* demonstrates that the observed cysts are not composed of GSCs. Scale bar for main images is 15 micron, scale bar for insets is 3.75 micron.

**Supplemental Figure 2.**
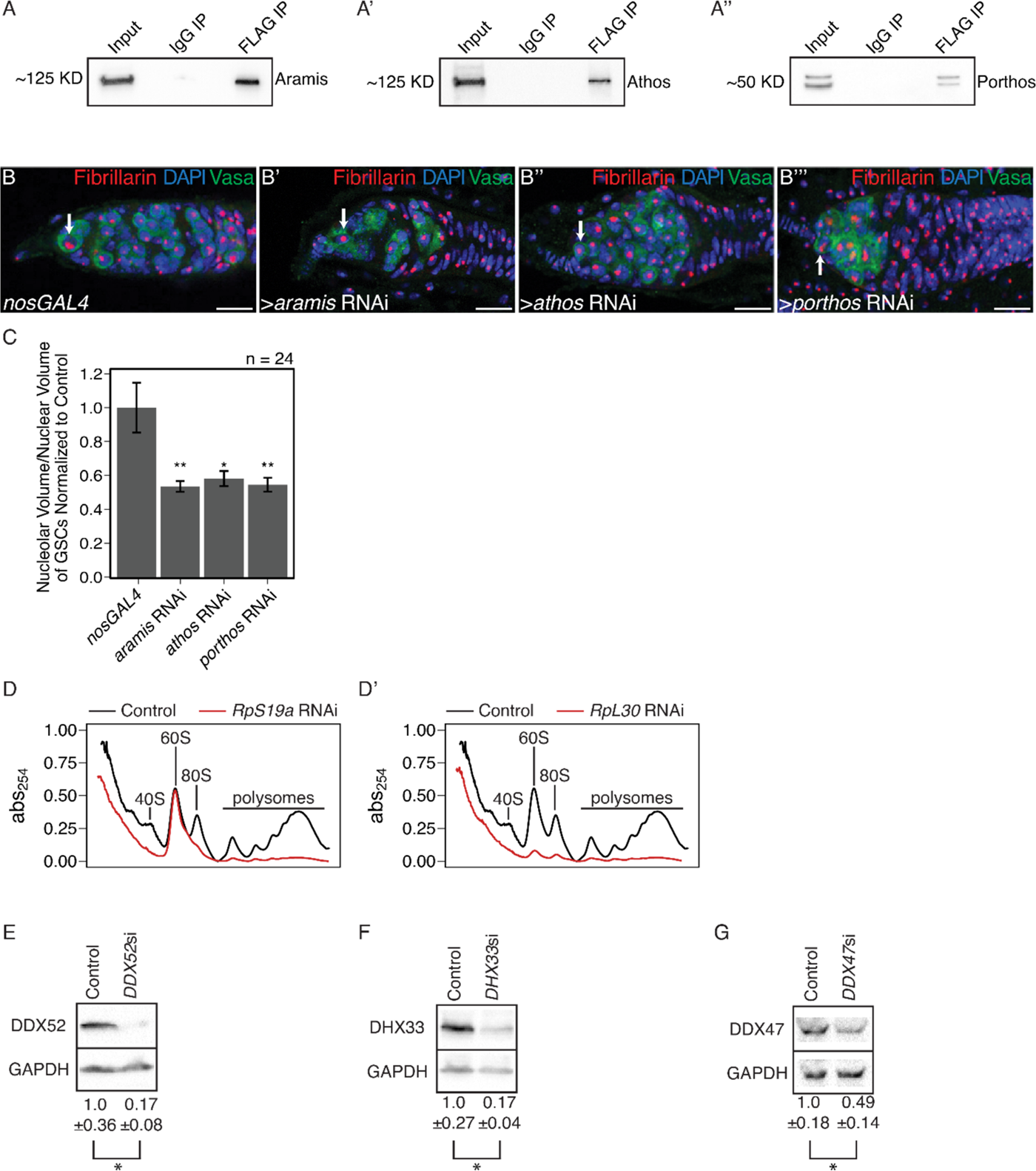
Athos, Aramis, and Porthos are required for efficient ribosome biogenesis., related to. **Figure 2**. (**A-A’’**) Western blots of immunoprecipitations from ovaries for FLAG-tagged (**A**) Aramis, (**A’**) Athos, and (**A’’**) Porthos. (**B-B’’’**) Confocal images of (**B**) *nosGAL4*, driver control, (**B’**) *aramis* (**B’’**) *athos* and (**B’’’**) *porthos* germline RNAi germaria stained for Fibrillarin (red), DAPI (blue), and Vasa (green). (**C**) Quantification of nucleolar volume in GSCs of *aramis*, *athos*, and *porthos* RNAi, compared to control normalized to somatic nucleolar volume indicates loss of each helicase results in nucleolar stress (n=24 GSCs per genotype, One-way ANOVA, p<0.001, with Welch’s t-test, * = p<0.05, ** = p<0.01). (**D-D’**) Polysome preparations from *Drosophila* S2 cells in cells treated with dsRNA targeting (**D**) *RpS19a* or (**D’**) *RpL30*. (**E-G**) Western blot against proteins targeted for depletion by siRNA in HeLa cells. The human homologs of (**E**) Aramis (DDX52), (**F**) Athos (DHX33), and (**G**) Porthos (DDX47) are efficiently depleted with siRNA treatment after 72 hours (n=3, Welch’s t-test, * = p<0.05). Scale bar for all images is 15 micron.

**Supplemental Figure 3.**
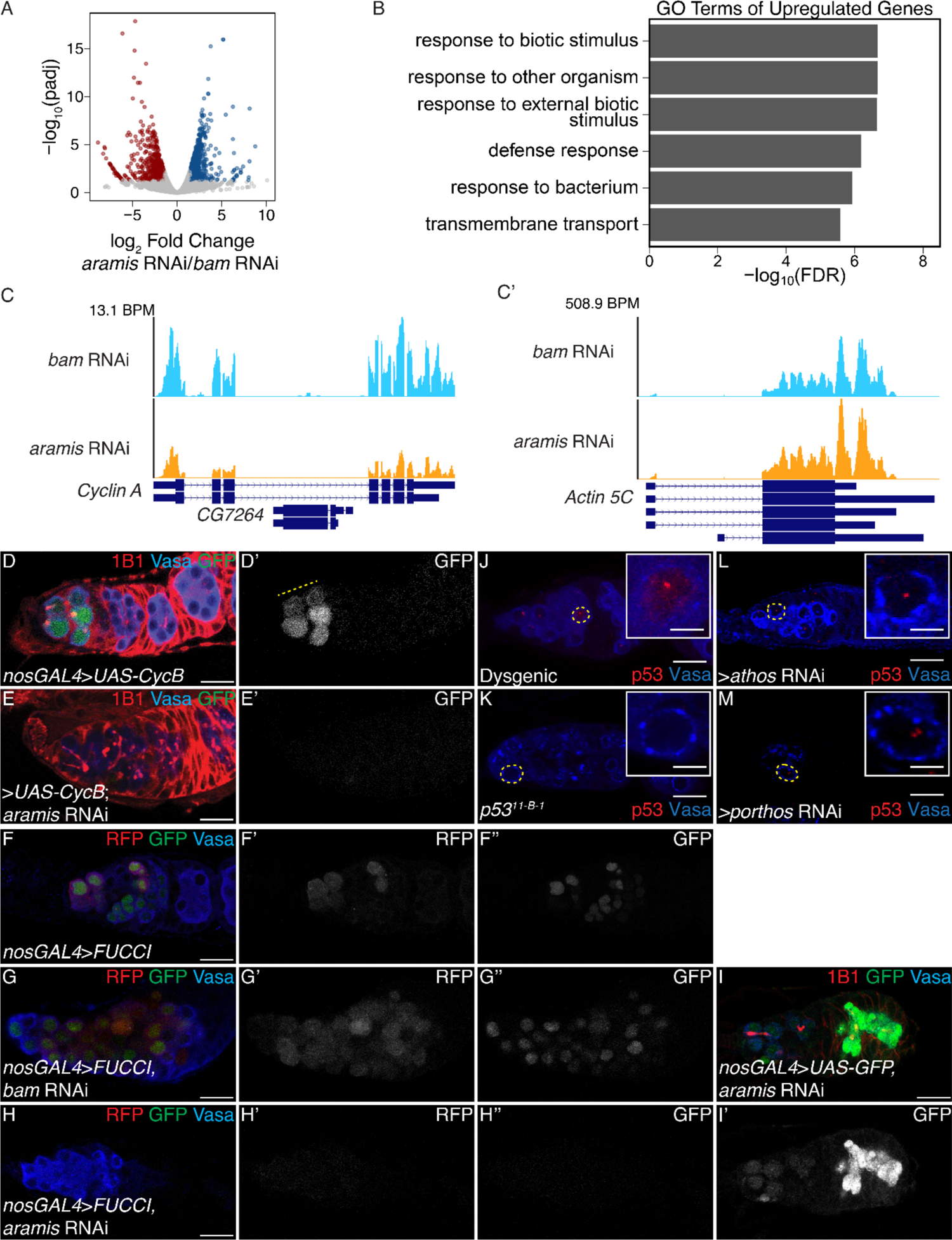
Aramis is required for proper cell cycle progression, related to. **Figure 3**. (**A**) Volcano plot of mRNA expression in *aramis* RNAi compared to *bam* RNAi. Blue points represent mRNAs significantly upregulated *aramis* RNAi compared to *bam* RNAi, red points represent mRNAs significantly downregulated *aramis* RNAi compared to *bam* RNAi. (**B**) Bar plot representing the most significant Biological Process GO terms of upregulated genes in ovaries depleted of *aramis* compared to the developmental control, *bam* RNAi. (**C-C’**) Genome browser tracks of mRNA expression at the (**C**) *Cyclin A* and (**C’**) *Actin 5C* loci indicate that the RNAseq target gene *Cyclin A* expression is downregulated, while a non-target, *Actin 5C* is not downregulated. (**D-E’**) Confocal images of germaria stained for 1B1 (red), Vasa (blue), and Cyclin B::GFP (green, grayscale) in (**D-D’**) control and (**E-E’**) germline depletion of *aramis* demonstrates that functional Cyclin B::GFP cannot be efficiently expressed in germline depleted of *aramis*. (**F-H’’**) Confocal images of germaria that express Fly-FUCCI in the germline stained for Vasa (blue). GFP-E2f1^degron^ (green, right grayscale) and RFP-CycB^degron^ (red, left grayscale) in (**F-F’’**) *nosGAL4*, driver control ovaries, (**G-G’’**) *bam* RNAi as a developmental control, and (**H-H’’**) ovaries with germline depletion *of aramis* demonstrates that the germline of *aramis* RNAi germline depleted ovaries are negative for both G1 and G2 cell cycle markers. (**I-I’**) Confocal images of *aramis* germline RNAi expressing GFP in the germline, stained for 1B1 (red), Vasa (blue), and GFP (green, grayscale) indicates productive translation of transgenes still occurs. (**J-M**) Confocal images of germaria stained for p53 (red) and Vasa (blue) in (**J**) hybrid dysgenic, Harwich, ovaries and (**K**) p53^11-B-1^ ovaries stained for p53 (red) and Vasa (blue) demonstrate the expected p53 staining patterns. (**L-M**) Confocal images of germaria stained for p53 (red) and Vasa (blue) in ovaries depleted of (**L**) *athos* or (**M**) *porthos* in the germline exhibit p53 punctate staining. Cells highlighted by a dashed yellow circle represent cells shown in the inset. Scale bar for main images is 15 micron, scale bar for insets is 3.75 micron.

**Supplemental Figure 4.**
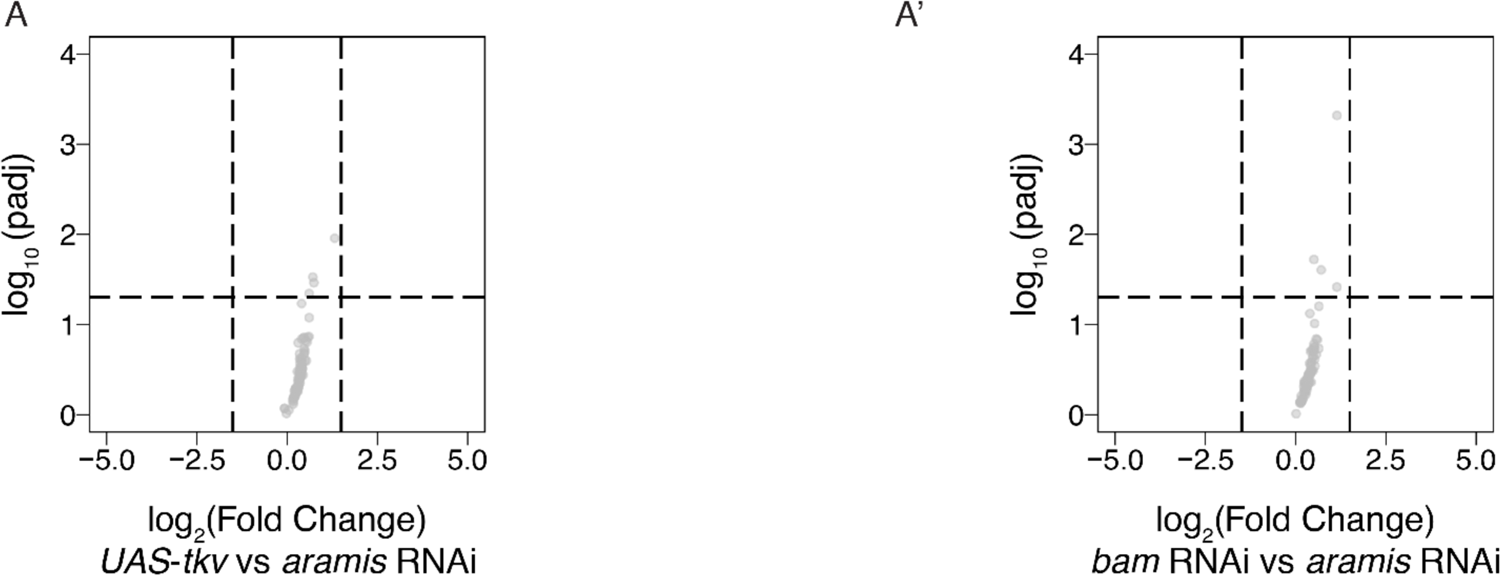
The mRNA levels of Aramis polysome-seq targets are not significantly changing, related to. **Figure 4**. (**A-A’**) Volcano plot of mRNA expression from poly(A)+ mRNA input libraries in germline *aramis* RNAi compared to (**A**) germline driven *UAS*-*tkv* and (**A’**) *bam* RNAi of targets identified from polysome-seq. No target genes identified from polysome-seq meet the differential expression cutoff for mRNA in *UAS-tkv* compared to *aramis* RNAi or *bam* RNAi compared to *aramis* RNAi input libraries.

**Supplemental Figure 5.**
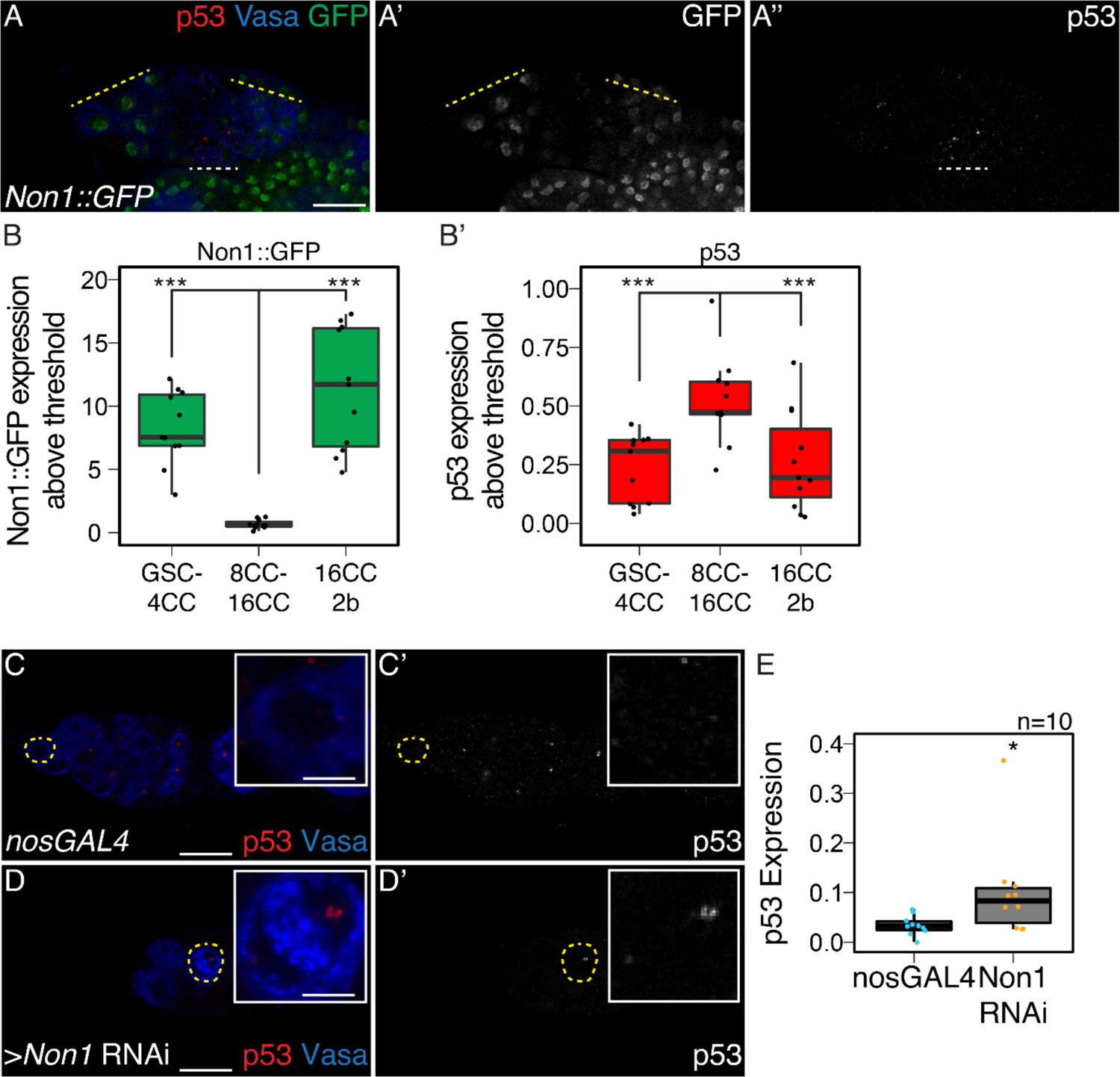
Non1 and p53 are inversely expressed, related to. **Figure 5**. (**A-A’’**) Confocal images of ovarioles expressing Non1::GFP stained for p53 (red, right grayscale), Vasa (blue), and Non1::GFP (green, left grayscale). (**B-B’**) Quantifications of staining, (**B**) peak Non1 expression in control ovaries occurs in GSC-4 cell cyst stages and 16-cell cyst-region 2b stages where (**B’**) p53 expression is low. (**C-D’**) Confocal images of *nosGAL4*, driver control (**C-C’**) and germline *Non1* RNAi germaria stained for p53 (red, grayscale) and Vasa (blue). (**E**) Quantification of p53 punctate area above cutoff are markedly brighter in the germline of *Non1* RNAi depleted ovaries compared to the control. Cells highlighted by a dashed yellow circle represent cells shown in the inset. Scale bar for main images is 15 micron, scale bar for insets is 3.75 micron.

**Supplemental Figure 6.**
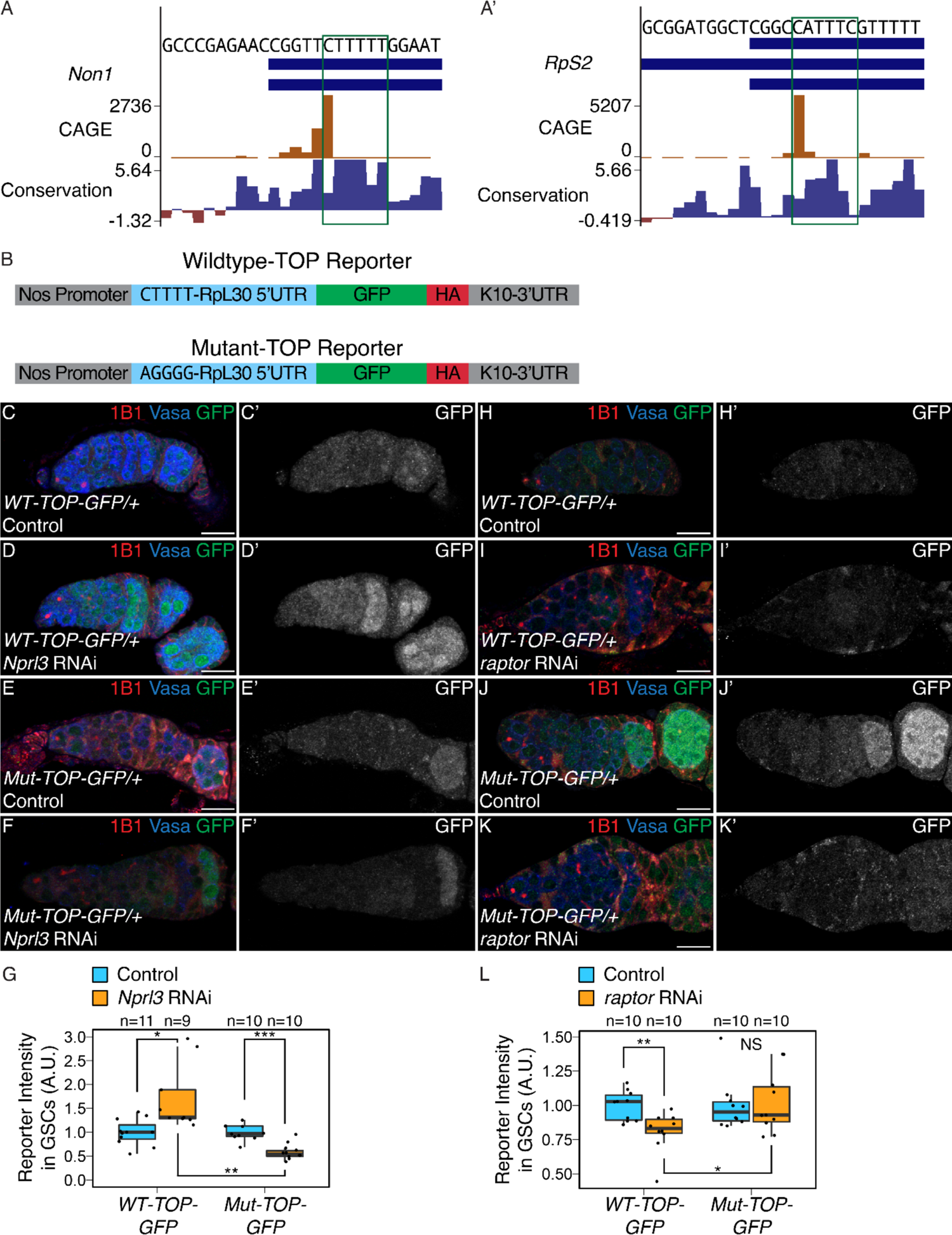
TORC1 activity regulates TOP expression in the germarium, related to. **Figure 6**. (**A-A’**) Genome browser tracks of the (**A**) *Non1* and (**A’**) *RpS2* loci in ovary CAGE-seq data showing the proportion of transcripts that are produced from a given TSS (orange). Predominant TSSs are shown in orange and putative TOP motif beginning at the dominant TSS is indicated with a green box. The bottom blue and red graph represents sequence conservation of the locus across *Diptera*. The dominant TSS of *Non1* initiates with a canonical TOP motif and the *RpS2* TSS initiates at a sequence resembling a TOP motif. (**B**) Diagram of the *WT* and *Mut-TOP-GFP* reporter constructs indicating the TOP sequence that is mutated by transversion in the Mutant reporter (blue). (**C-D’**) Confocal images of *WT-TOP* reporter expression stained for 1B1 (red), GFP (green, grayscale), and Vasa (blue) in (**C-C’**) *nosGAL4*, driver control ovaries and (**D-D’**) ovaries depleted of *Nprl3* in the germline. (**E-F’**) Confocal images of *Mut-TOP-GFP* reporter expression stained for 1B1 (red), GFP (green, grayscale), and Vasa (blue) in (**E-E’**) *nosGAL4*, driver control ovaries and (**F-F’**) ovaries depleted of *Nprl3* in the germline. (**G**) A.U. quantification of WT and Mutant TOP reporter expression in GSCs of *nosGAL4*, driver control ovaries and GSCs of *Nprl3* germline depleted ovaries normalized to Vasa expression indicate that the relative expression of the *WT-TOP-GFP* reporter is higher than the *Mut-TOP-GFP* reporter (n=9-11 germaria per genotype, Welch’s t-test, * = p<0.05, ** = p<0.01, *** = p<0.001). (**H-I’**) Confocal images of *WT-TOP* reporter expression stained for 1B1 (red), GFP (green, grayscale), and Vasa (blue) in (**H-H’**) *nosGAL4*, driver control ovaries and (**I-I’**) ovaries depleted of *raptor* in the germline. (**J-K’**) Confocal images of *Mut-TOP-GFP* reporter expression stained for 1B1 (red), GFP (green, grayscale), and Vasa (blue) in (**J-J’**) *nosGAL4*, driver control ovaries and (**K-K’**) ovaries depleted of *raptor* in the germline. (**L**) A.U. quantification of WT and Mutant TOP reporter expression in GSCs of *nosGAL4*, driver control ovaries and GSCs of *raptor* germline depleted ovaries normalized to Vasa expression indicate that the relative expression of the *WT-TOP-GFP* reporter is lower than the *Mut-TOP-GFP* reporter (n=10 germaria per genotype, Welch’s t-test, * = p<0.05, ** = p<0.01). Scale bar for images is 15 micron.

**Supplemental Figure 7.**
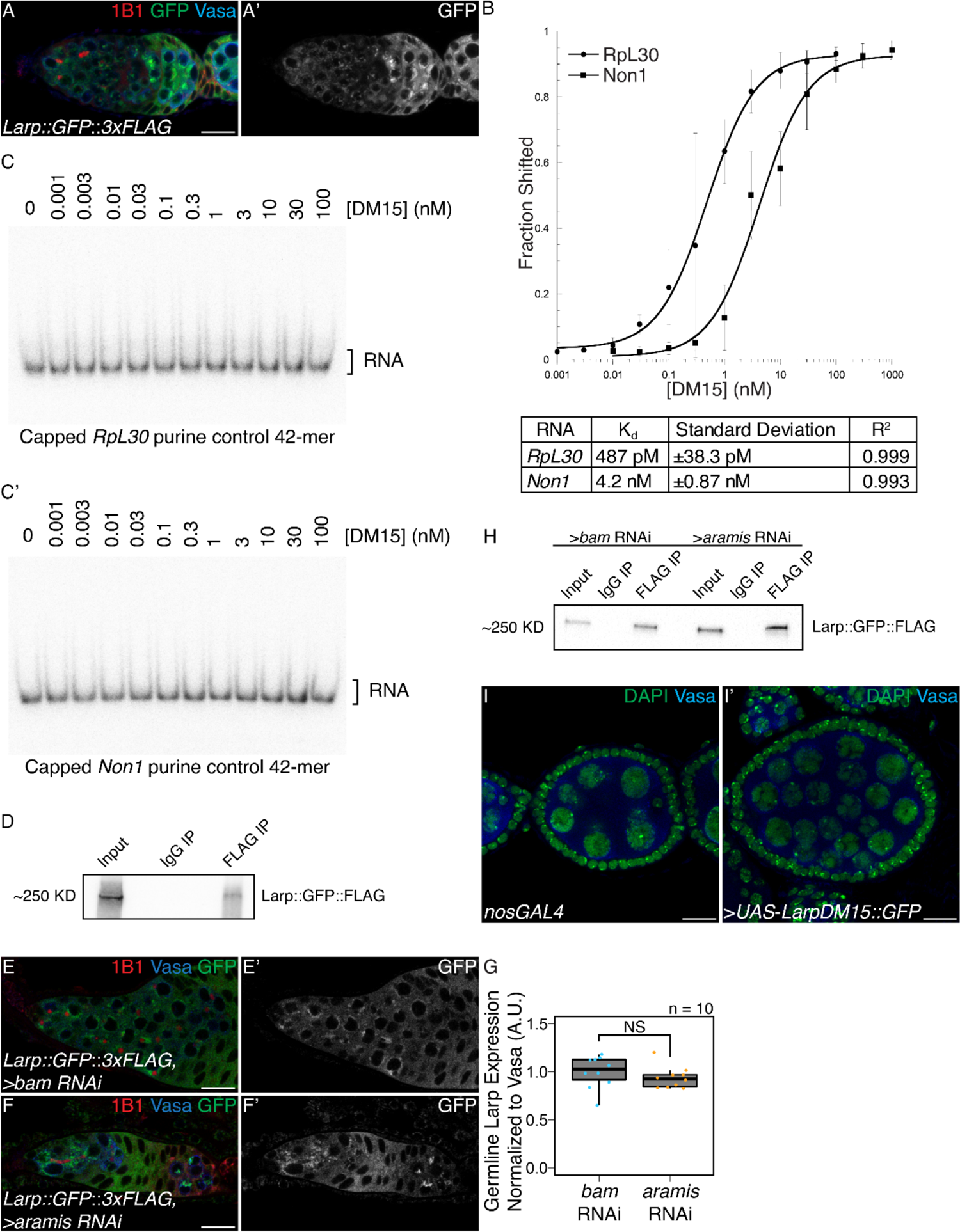
Larp binds specifically to TOP containing mRNAs and regulates cytokinesis, related to. **Figure 7**. (**A-A’**) Confocal images of germaria stained for 1B1 (red), Vasa (blue), and *Larp GFP-3xFLAG* (green, grayscale) indicates Larp is expressed throughout early oogenesis. (**B**) Quantification of EMSAs and summary of K_d_ of the protein-RNA interactions. (**C-C’**) EMSA of Larp-DM15 and the leading 42 nucleotides of (**B**) *RpL30* and (**B’**) *Non1* with their TOP sequence mutated to purines as a negative control with increasing concentrations of Larp-DM15 from left to right indicates that Larp-DM15 requires a leading TOP sequence for its binding. (**D**) Western of representative IP of Larp::GFP::FLAG from ovary tissue used for RNA IP-seq. (**E-F’**) Confocal images of *Larp::GFP::FLAG* reporter expression stained for 1B1 (red), GFP (green, grayscale), and Vasa (blue) in (**E-E’**) *bam* and (**F-F’**) *aramis* depleted germaria. (**G**) A.U. quantification of Larp::GFP::FLAG reporter expression in the germline of *bam* RNAi and *aramis* RNAi demonstrates that the germline expression of Larp is not elevated in *aramis* germline RNAi compared to *bam* germline RNAi as a developmental control (n=10, p>0.05, Welch’s t-test). (**H**) Western of representative IP of Larp::GFP::FLAG from ovary tissue used for RNA IP qPCR. (**I-I’**) Confocal images of (**I**) *nosGAL4*, driver control and (**I’**) ovaries overexpressing the DM15 region of Larp in the germline ovaries stained for DAPI (green) and Vasa (blue). Overexpression of Larp-DM15 results in the production of 32-cell egg chambers which indicates it causes a cytokinesis defect. Scale bar for all images is 15 micron.

**Supplemental Table 1. Results of germline helicase RNAi screen on ovariole morphology.** Results of screen of RNA helicases depleted from the germline. Reported is the majority phenotype from n=50 ovarioles.

**Supplemental Table 2. Differential expression analysis from RNAseq of ovaries depleted of *aramis* in the germline compared to a developmental control**. DEseq2 output from RNAseq of ovaries depleted of *aramis* in the germline compared to ovaries depleted of bam in the germline as a developmental control. Sheet 1 (Downregulated Genes) contains genes and corresponding DEseq2 output meeting the cutoffs to be considered downregulated in aramis RNAi compared to bam RNAi. Sheet 2 (Upregulated Genes) contains genes and corresponding DEseq2 output meeting the cutoffs to be considered upregulated in aramis RNAi compared to bam RNAi. Sheet 3 (All Genes) contains DEseq2 output for all genes in the dm6 assembly.

**Supplemental Table 3. Analysis of polysome-seq of ovaries depleted of *aramis* in the germline compared to developmental controls**. Results of polysome-seq from ovaries depleted of *aramis* in the germline, ovaries depleted of *bam*, and ovaries overexpressing tkv in the germline as developmental controls. Sheet 1 (Downregulated Genes) contains genes and corresponding polysome/input ratio values and values representing the difference in the polysome/input ratios between *aramis* RNAi and the developmental controls meeting the cutoffs to be considered downregulated in *aramis* RNAi. Sheet 2 (Upregulated Genes) contains genes and corresponding polysome/input ratio values and values representing the difference in the polysome/input ratios between *aramis* RNAi and the developmental controls meeting the cutoffs to be considered upregulated in *aramis* RNAi. Sheet 3 (All Genes) contains DEseq2 output for all genes in the dm6 assembly.

**Supplemental Table 4. Aramis translation targets contain TOP sequences.** List of *aramis* RNAi polysome downregulated targets and the position and sequence of the first instance of a 5-mer pyrimidine sequence downstream of the CAGE-defined TSS of each gene.

**Supplemental Table 5. Enrichment analysis of Larp RNA IP mRNA-seq**. Results of Larp::GFP::FLAG IP/IgG/Input mRNAseq. Each sheet contains the output of DEseq2. Sheet 1 (Larp Targets) contains Larp IP targets as defined in methods. Sheet 2 (IP vs In Enriched) contains genes significantly enriched in the Larp IP samples compared to the input samples. Sheet 3 (IgG vs In Enriched) contains genes significantly enriched (see methods) in the IgG samples compared to the input samples. Sheet 4 (IPvsIn All Genes) contains the DEseq2 output of all genes in the Larp IP samples compared to the input samples. Sheet 5 (IgG vs In All Genes) contains the DEseq2 output of all genes in the IgG samples compared to the input samples.

## Materials and Methods

### Fly lines

The following Bloomington Stock Center lines were used in this study: #25751 *UAS-Dcr2;nosGAL4*, #4442 *nosGAL4;MKRS/TM6*, #32334 Aramis RNAi#1 CG5589^HMS00325^, #56977 Athos RNAi#1 CG4901^HMC04417^, #36589 Porthos RNAi#1 CG9253^GL00549^, #36537 UAS-tkv.CA, #33631 bam RNAi^HMS00029^, #6815 p53^5A-1-4^, #4264 Harwich, #6816 p53^11-1B-1^, #55101 FUCCI: UASp-GFP.E2f1.1-230, UASp-mRFP1.CycB.1-266/TM6B, #5431 UAS-EGFP, #78777 Non1 RNAi P{TRiP.HMS05872}, #61790 Larp::GFP::3xFLAG Mi{PT-GFSTF.1}larp^MI06928-GFSTF.1^, #8841 w[1118]; Df(3R)Hsp70A, Df(3R)Hsp70B, #55384 Nprl3 RNAi P{TRiP.HMC04072}attP40, #34814 raptor RNAi P{TRiP.HMS00124}attP2 The following Vienna Stock Center lines were used in this study: Aramis RNAi#2 CG5589^v44322^, Athos RNAi#2 CG4901^v34905^, Aramis::GFP PBac{fTRG01033.sfGFP-TVPTBF}VK00002, Athos::GFP PBac{fTRG01233.sfGFP-TVPTBF}VK00033, Non1::GFP PBac{fTRG00617.sfGFP-TVPTBF}VK00033 The following additional fly lines were used in the study: UASp-CycB::GFP (Mathieu et al., 2013), *UAS-Dcr2;nosGAL4;bamGFP, If/CyO;nosGAL4* (Lehmann Lab), w1118 (Lehmann lab), *tjGAL4/CyO* (Tanentzapf et al., 2007), RpS2::GFP^CB02294^ (Buszczak et al., 2007; Zhang et al., 2014), UASp-Non1 (this study), UASp-Larp-DM15 (this study), WT-TOP-Reporter (this study), Mutant-TOP-Reporter (this study).

### Antibodies IF

The following antibodies were used for immunoflourenscence: mouse anti-1B1 1:20 (DSHB 1B1), rabbit anti-Vasa 1:833-1:4000 (Rangan Lab), chicken anti-Vasa 1:833-1:4000 (Rangan Lab) (Upadhyay et al., 2016), rabbit anti-pTyr 1:500 (Sigma T1235), rabbit anti-pMad 1:200 (Abcam ab52903), rabbit anti-GFP 1:2000 (abcam, ab6556), mouse anti-p53 1:200 (DSHB 25F4), Rabbit anti-CycB 1:200 (Santa Cruz Biotechnology, 25764), Rabbit anti-Fibrillarin 1:200 (Abcam ab5821), Mouse anti-Fibrillarin 1:50 (Fuchs Lab) (McCarthy et al., 2018). Alexa 488 (Molecular Probes), Cy3 and Cy5 (Jackson Labs) were used at a dilution of 1:500.

### Antibodies Western/IP

Mouse anti-FLAG-HRP 1:5000 (Sigma Aldrich, A8592) Mouse anti-FLAG (Sigma Aldrich, F1804) Anti-GAPDH-HRP 1:10,000 (Cell Signaling, 14C10) Rabbit anti-DDX52 1:5000 (Bethyl, A303-053A) Rabbit anti-DHX33 1:5000 (Bethyl, A300-800A) Rabbit anti-DDX47 1:1000 (Bethyl, A302-977A)

### Protein Domain Analysis

Protein domain figures were adapted from: The Pfam protein families database in 2019: S. El-Gebali et al. Nucleic Acids Research (2019). Protein Similarity values were obtained from the DRSC/TRiP Functional Genomics Resources.

### TOP Reporter Cloning

gBlocks (see primer list for details) were cloned into pCasper2 containing a Nos promoter, HA-tag, GFP-tag, and K10 3’UTR. PCR was used in order to amplify the gBlock and to remove the 5’-end of the RpL30 5’UTR in order to generate the 5’-UTR discovered via CAGE-seq. In order to clone the Nos promoter followed by the RpL30 5’UTR without an intervening restriction site, the portion of the plasmid 5’ of the 5’UTR consisting of a portion of the plasmid backbone, a NotI restriction site, and the Nos Promoter was amplified from the pCasper plasmid using PCR. HiFi cloning was performed on the amplified fragments. The backbone was cut with NotI and SpeI and HiFi cloning was performed according to the manufactures’ instructions except the HiFi incubation was performed for 1 hour to increase cloning efficiency. Colonies were picked and cultured and plasmids were purified using standard techniques. Sequencing was performed by Eton Bioscience Inc. to confirm the correct sequence was present in the final plasmids. Midi-prep scale plasmid was prepared using standard methods and plasmids were sent to BestGene Inc. for microinjection.

### Gateway Cloning

Gateway cloning was performed as described according to the manufacture’s manual. Briefly, primers containing the appropriate Gateway *attb* sequence on the 5’-ends and gene specific sequences on the 3’-ends (see primer list for sequences) were used to PCR amplify each gene of interest. PCR fragments were BP cloned into pEntr221 as detailed in the Thermofisher Gateway Cloning Manual and used to transform Invitrogen One Shot OmniMAX 2 T1 Phage-Resistant Cells. Resulting clones were picked and used to perform LR cloning into either pPGW or pPWG as appropriate. Cloning was carried out according to the Thermofisher Gateway Cloning Manual except the LR incubation was carried out up to 16 hours. Colonies were picked and cultured and plasmids were purified using standard techniques. Sequencing was performed by Eton Bioscience Inc. to confirm the correct sequence was present in the final plasmids. Midi-prep scale plasmid was prepared using standard methods and plasmids were sent to BestGene Inc. for microinjection.

### Egg Laying Test

Newly eclosed flies were collected and fattened overnight on yeast. Six female flies were crossed to 4 male controls and kept in cages at 25°C. Flies were allowed to lay for three days, and plates were changed and counted daily. Total number of eggs laid over the three day laying periods were determined and averaged between three replicate crosses for control and experimental crosses.

### Immunostaining

Ovaries were dissected and teased apart with mounting needles in cold PBS and kept on ice for subsequent dissections. All incubations were performed with nutation. Ovaries were fixed for 10-15 min in 5% methanol-free formaldehyde in PBS. Ovaries were washed with PBT (1x PBS, 0.5% Triton X-100, 0.3% BSA) once quickly, twice for 5 min, and finally for 15 min. Ovaries were incubated overnight, up to 72 hours in PBT with the appropriate primary antibodies. Ovaries were again washed with PBT once quickly, twice for 5 min, and finally for 15 min. Ovaries were then incubated with the appropriate secondary antibodies in PBT overnight up to 72 hours at 4°C. Ovaries were washed once quickly, twice for 5 min, and finally for 15 min in PBST (1x PBS, 0.2% Tween 20 Ovaries). Ovaries were mounted with Vectashield with 4′,6-diamidino-2-phenylindole (DAPI) (Vector Laboratories) and imaged on a Zeiss 710. All gain, laser power, and other relevant settings were kept constant for any immunostainings being compared. Image processing was performed in Fiji, gain was adjusted, and images were cropped in Photoshop CC 2018.

### Florescent Imaging

Tissues were visualized and imaged were acquired using a Zeiss LSM-710 confocal microscope under the 20× and 40× oil objectives.

### Measurement of global protein synthesis

OPP (Thermo Fisher, C10456) treatment was performed as in McCarthy (2019). Briefly, ovaries were dissected in Schneider’s media (Thermo Fisher, 21720024) and incubated in 50 μM of OPP reagent for 30 minutes. Tissue was washed in 1x PBS and fixed for 10 minutes in 1x PBS plus 5% methanol-free formaldehyde. Tissue was permeabilized with 1% Triton X-100 in 1x PBST (1x PBS, 0.2% Tween 20) for 30 minutes. Samples were washed with 1x PBS and incubated with Click-iT reaction cocktail, washed with Click-iT reaction rinse buffer according to manufacturer’s instructions. Samples were then immunostained according to previously described procedures.

### Image Quantifications

All quantifications were performed on images using the same confocal settings. A.U. quantifications were performed in Fiji on images taken with identical settings using the “Measure” function. Intensities were normalized as indicated in the figure legends, boxplots of A.U. measurements were plotted using R and statistics were calculated using R. Quantification of nucleolar size was measured in Fiji by measuring the diameter of the nucleolus using the measure tool in Fiji. Volumes were calculated using the formula for a sphere. Quantification of p53 area of expression was performed from control, *nosGAL4* and nosGAL4>*aramis* RNAi germaria. A manual threshold was set based off of qualitative assessment of a “punctate”. For control ovaries, cells proximal to the niche consisting of GSCs/CBs were outlined and for *aramis* RNAi the entire germline proximal to the niche was outlined and a Fiji script was used to determine the number of pixels above the threshold and the total number of pixels. Data from each slice for each replicate was summed prior to plotting and statistical analysis.

Colocalization analysis of helicases with Fibrillarin was performed in Fiji using the Plot Profile tool. A selection box was drawn over a Fibrillarin punctate of interest (indicated with a box in the images) and Plot Profiles was acquired for each channel of interest. Data was plotted and Spearman correlations calculated using R.

Quantification of Non1-GFP expression and p53 expression over development was calculated in Fiji using the Auto Threshold tool with the Yen method (Sezgin and Sankur, 2004) to threshold expression. Quantifications were performed on 3 merged slices and egg chambers were cropped out of quantified images prior to thresholding to prevent areas outside of the germarium from influencing the thresholding algorithm. Areas of germline with “high” and “low” expression of Non1-GFP were outlined manually and a custom Fiji script was used in order to quantify the proportion of pixels in the selected marked as positive for expression for either Non1-GFP or p53, staging was inferred from the results of the Non1-GFP quantification performed using 1B1 to determine the stages of peak Non1 expression. Percent area was plotted with ggplot2 as boxplots in a custom R script.

### RNA Extraction from Ovaries

RNA extraction was performed using standard methods. Ovaries were dissected into PBS and transferred to microcentrifuge tubes. PBS was removed and 100ul of Trizol was added and ovaries were flash frozen and stored at −80 °C. Ovaries were lysed in the microcentrifuge tube using a plastic disposable pestle. Trizol was added to 1 mL total volume and sample was vigorously shaken and incubated for 5 min at RT. The samples were centrifuged for x min at >13,000 g at 4 °C and the supernatant was transferred to a fresh microcentrifuge tube. 500 ul of chloroform was added and the samples were vigorously shaken and incubated for 5 minutes at RT. Samples were spun at max speed for 10 minutes at 4 °C. The supernatant was transferred to a fresh microcentrifuge tube and ethanol precipitated. Sodium acetate was added equaling 10% of the volume transferred and 2-2.5 volumes of 100% ethanol were added. The samples were shaken thoroughly and left to precipitate at −20 °C overnight. The samples were centrifuged at max speed at 4 °C for 15 min to pellet the RNA. The supernatant was discarded and 500 ul of 75% ethanol was added to wash the pellet. The samples were vortexed to dislodge the pellet to ensure thorough washing. The samples were spun at 4 °C for 5 min and the supernatant was discarded. The pellets were left for 10-20 min until dry. The pellets were resuspended in 20-50ul of RNAse free water and the absorbance at 260 was measured on a nanodrop to measure the concentration of each sample.

### S2 Cell RNAi

DRSC-S2 cells (Stock #181, DGRC) were cultured according to standard methods in M3+BPYE media supplemented with 10% heat-inactivated FBS. dsRNA for RNAi was prepared as described by the SnapDragon manual. Briefly, template was prepared from S2 cell cDNA using the appropriate primers (see primer list) designed using SnapDragon (https://www.flyrnai.org/snapdragon). For *in-vitro* transcription the T7 Megascript kit (AM1334) was used following manufacturer’s instructions and in-vitro transcriptions were incubated overnight at 37°C. The RNA was treated with DNAse according to the T7 Megascript manual and the RNA was purified using acid-phenol chloroform extraction and ethanol precipitated. The resulting RNA was annealed by heating at 65°C for 5 minutes and slow cooling to 37°C for an hour. S2 cell RNAi was performed essentially as previously described using Effectine (Zhou et al., 2013). 1.0×10^6^ cells were seeded 30 minutes prior to transfection and allowed to attach. After 30 minutes, just prior to transfection, the media was changed for 500 µl of fresh media. 500 µl of transfection complexes using 1 µg of dsRNA was prepared per well of a 6-well plate and pipetted dropwise onto seeded cells. After 24 hours an additional 1 mL of media was added to each well. After an additional 24 hours cells were passaged to 10 cm dishes. After an additional 3 days cells were harvested for further analysis.

### HeLa Cell RNAi

HeLa cells were cultured under standard conditions in DMEM (Gibco) supplemented with 10% FBS, and 2 mM L-glutamine at 37°C and 5% CO_2_. RNAi was performed in HeLa cells using the siRNAs in the primer list. 5% of cells from a 10 cm dishes with cells between 70-85% confluency were seeded to 6-well plates. Transfection was performed the following day using Dharmafect (PerkinElmer T-2001-03) according to the manufacture’s procedure. Briefly, per well to be transfected Transfection Master Mix was prepared by mixing 5 µl of Dharmafect with 200 µl of OptiMEM (ThermoFisher 31985070) and incubated for 5 minutes at RT. 50 µl of OptiMEM was mixed with the given 3.75 µl of 20 µM siRNA and added to 100 µl of Transfection Master Mix and incubated for 20-30 minutes at RT. Each transfection reaction was added to 600 µl of DMEM. Media from the previous day was removed from the cells and replaced with the transfection mix. Cells were incubated at 37°C for one day, then passaged to 10 cm dishes and cultured for an additional two days. Cells were subsequently prepared for Western Blotting or Polysome Profiling (see respective sections).

### Polysome-profiling

Polysome-profiling in S2 and HeLa cells was performed as in Fuchs et al. (2011) with minor modifications. HeLa cells were washed in cold PBS and lysed on the plate by scraping under 400 µl lysis buffer (300 mM NaCl, 15 mM Tris-HCl, pH 7.5, 15 mM EDTA, 100 μg/mL cycloheximide, 1% Triton X-100). S2 cells were resuspended by pipetting, pelleted by centrifugation at 800g for one minute, and washed in cold PBS. S2 cells were again pelleted and resuspended in 400 µl of lysis buffer. HeLa and S2 cells were then allowed to continue to lyse for 15 min on ice. Lysate was cleared by centrifugation at 8500g for 5 min at 4°C. Cleared lysate was loaded onto 10%-50% sucrose gradients (300 mM NaCl, 15 mM Tris-HCl, pH 7.5, 15 mM MgCl2, 100 g/mL cycloheximide) and centrifuged in an SW41 rotor at 35,000 RPM, for 3 hours. Gradients were fractionated on a Density Gradient Fractionation System (Brandel, #621140007) at 0.75 mL/min. Data generated from gradients were plotted using R.

### Western Blot

HeLa cells were harvested for Western by in RIPA buffer by scraping. Western blotting were performed according to standard methods, briefly, each sample was loaded onto a 4-20% commercial, precast gels and run at 100V for 60-90m depending on the size of the protein of interest. Gels were transferred to nitrocellulose membranes at 100V for 1hr at 4°C. Blot was blocked in 1% milk in PBS and washed 3 times with PBS-T for 5 minutes. Primary antibodies were diluted in PBS-T+5% BSA and incubated overnight. Blot was washed once quickly, once for 5m, and once for 10m in PBS-T. Blot was subsequently imaged with ECL for conjugated primaries. For unconjugated primaries, the appropriate secondary was diluted 1:10,000 in 5% milk and incubated for 2-4 hours at RT. Blot was washed once quickly, once for 5m, and once for 10m in PBS-T and imaged. Images were quantified using Fiji.

### mRNAseq Library Preparation and Analysis

Libraries were prepared with the Biooscientific kit (Bioo Scientific Corp., NOVA-5138-08) according to manufacturer’s instructions with minor modifications. Briefly, RNA was prepared with Turbo DNAse according to manufacturer’s instructions (TURBO DNA-free Kit, Life Technologies, AM1907), and incubated at 37°C for 30 min. DNAse was inactivated using the included DNAse Inactivation reagent and buffer according to manufactures instructions. The RNA was centrifuged at 1000 g for 1.5 min and 19 μl of supernatant was transferred into a new 1.5 mL tube. This tube was again centrifuged at 1000 g for 1.5 min and 18 μl of supernatant was transferred to a new tube to minimize any Inactivation reagent carry-over. RNA concentration was measured on a nanodrop. Poly-A selection was performed on a normalized quantity of RNA dependent on the lowest amount of RNA in a sample, but within the manufacturer’s specifications for starting material. Poly-A selection was performed according to manufacturer’s instructions (Bioo Scientific Corp., 710 NOVA-512991). Following Poly-A selection mRNA libraries were generated according to manufactures instructions (Bioo Scientific Corp., NOVA-5138-08) except RNA was incubated for 13 min at 95°C to generate optimal fragment sizes. Library quantity was assessed via Qubit according to manufacturer’s instructions and library quality was assessed with a Bioanalyzer or Fragment Analyzer according to manufacturer’s instructions to assess the library size distribution. Sequencing was performed on biological duplicates from each genotype on an Illumina NextSeq500 by the Center for Functional Genomics (CFG) to generate single end 75 base pair reads. Reads were aligned to the dm6.01 assembly of the Drosophila genome using HISAT v2.1.0. Reads were counted using featureCounts v1.4.6.p5. UCSC genome browser tracks were generated using the bam coverage module of deeptools v3.1.2.0.0. Differential expression analysis was performed using DEseq2 (v1.24.0) and data was plotted using R. Differentially expressed genes were those with log_2_(foldchange) > |1.5| and FDR < 0.05. GO-term analysis of GO biological processes was performed on differentially expressed genes using PANTHER via http://geneontology.org/. Fisher’s exact test was used to calculate significance and FDR was used to correct for multiple testing. GO-term analysis results were plotted using R.

### Polysome-seq

Polysome-seq was performed as in Flora et al. (2018b) with minor modifications. Ovaries were dissected in PBS and transferred to a microcentrifuge tube in liquid nitrogen. Ovaries were lysed in 300 µl of lysis buffer (300 mM NaCl, 15 mM Tris-HCl, pH 7.5, 15 mM EDTA, 100 μg/mL cycloheximide, 1% Triton X-100) and allowed to lyse for 15 min on ice. Lysate was cleared by centrifugation at 8500g for 5 min at 4°C. 20% of the lysate was reserved as input, 1 mL of Trizol (Invitrogen, 15596026) was added and RNA was stored at −80°C. Cleared lysate was loaded onto 10%-50% sucrose gradients (300 mM NaCl, 15 mM Tris-HCl, pH 7.5, 15 mM MgCl2, 100 g/mL cycloheximide) and centrifuged in an SW41 rotor at 35,000 RPM, for 3 hours. Gradients were fractionated on a Density Gradient Fractionation System (Brandel, #621140007) at 0.75 mL/min, 20 μl of 20% SDS, 8 μl of 0.5 M pH 8 EDTA, and 16 μl of proteinase K (NEB, P8107S) was added to each polysome fraction. Fractions were incubated for 30m at 37°C. Standard acid phenol chloroform purification followed by ethanol precipitation was performed on each fraction. The RNA from polysome fractions was pooled and RNAseq libraries were prepared.

### S2 Polysome-seq Data Analysis

Reads were checked for quality using FastQC. Reads were mapped to the *Drosophila* genome (dm6.01) using Hisat version 2.1.0. Mapped reads were assigned to features using featureCount version v1.6.4. Translation efficiency was calculated as in (Flora et al., 2018) using a custom R script. Briefly, CPMs (counts per million) values were calculated. Any gene having zero reads in any library was discarded from further analysis. The log_2_ ratio of CPMs between the polysome fraction and total mRNA was calculated and averaged between replicates. This ratio represents TE, TE of each replicate was averaged and delta TE was calculated (Porthos RNAi TE)/(GFP RNAi TE). This ratio represents delta TE. The group of highly expressed targets was defined as transcripts falling greater or less than one standard deviation from the median of delta TE with log_2_(TPM) expression greater than five.

### CAGE-seq Tracks

CAGE-seq tracks were visualized using the UCSC Genome Browser after adding the publicly available track hub ‘EPD Viewer Hub’.

### CAGE-seq Data Reanalysis

Publicly available genome browser tracks were obtained of CAGE-seq data (generated by Chen et al. (2014) and viewed through the UCSC Genome Browser. The original CAGE-seq data from ovaries was obtained from SRA under the accession number SRR488282. Reads were aligned to the dm6.01 assembly of the *Drosophila* genome using HISAT v2.1.0. cageFightR was used to determine the dominant TSS for every gene with sufficient expression in from the aligned dataset according to its documentation with default parameters excepting the following: For getCTSS, a mappingQualityThreshold of 10 was used. For normalizeTagCount the method used was “simpleTPM”. For clusterCTSS the following parameters were used; threshold = 1, thresholdIsTpm = TRUE, nrPassThreshold = 1, method = “paraclu”, maxDist = 20, removeSingletons = TRUE, keepSingletonsAbove = 5. Custom Rcripts were used in order to obtain genome sequence information downstream of the TSS of each gene identified.

### Motif Enrichment Analysis

Motif enrichment analysis was performed using Homer (Heinz et al., 2010) using the findmotifs.pl module, supplying Homer with the first 200 nucleotides downstream of the TSS as determined by CAGE-seq for polysome-seq targets and non-targets as a background control with the following parameters “-rna -nogo -p 6 -len 6”. Only motifs not marked as potential false positives were considered. The position of the putative TOP motifs was determined using a custom R script by searching for the first instance of any five pyrimidines in a row within the first 200 nucleotides of the TSS using the Biostrings package (Pagès et al., 2019). Results were plotted as a histogram in R.

### RNA Immunoprecipitation (RNA IP)

All RIPs were performed with biological triplicates. 50-60 ovary pairs were dissected for each sample in RNase free PBS and dissected ovaries were kept on ice during subsequent dissections. After dissection, ovaries were washed with 500 µl of PBS to remove any debris. This PBS was removed, and ovaries were lysed in 100 µl of RIPA buffer (10 mM Tris-Cl Buffer (pH 8.0), 1 mM EDTA, 1% Triton X-100,0.1% Sodium deoxycholate, 0.1% SDS, 140 mM NaCl, 1 mM PMSF, 1 cOmplete, EDTA-free Protease Inhibitor/10mLbuffer (Roche, 11873580001), RNase free H2O) supplemented with 8 µl of RNase Out. Following lysis an additional 180 µl of RIPA was added to each sample. Lysate was cleared with centrifugation at 14,000g for 20m at 4°C. Cleared lysate was transferred to a new 1.5 mL tube. 10% of this lysate was reserved for RNA input and 5% was reserved as a protein input. To the RNA input 100 µl of Trizol was added and the input was stored at −80°C. To the protein input SDS loading buffer was added to a 1X working concentration and the sample was heated at 95°C for 5m and stored at −20°C. The remaining lysate was equally divided into two new 1.5 mL tubes. To one tube 3 µg of mouse anti-FLAG antibody was added and to the other tube 3 µg of mouse IgG was added. These samples were incubated for 3 hours with nutation at 4°C. NP40 buffer was diluted to a 1X working concentration from a 10X stock (10x NP40 Buffer: 50 mM Tris-Cl Buffer (pH 8.0), 150 mM NaCl, 10% NP-40, 1 cOmplete, EDTA-free Protease Inhibitor Cocktail Pill/10mL buffer, RNase free H2O). 30 µl of Protein-G beads per RIP were pelleted on a magnetic stand and supernatant was discarded. 500 µl of 1X NP40 buffer was used to resuspend Protein-G beads by nutation. Once beads were resuspended, they were again pelleted on the magnetic stand. This washing process was repeated a total of 5 times. Washed Protein-G beads were added to each lysate and incubated overnight. The next day fresh 1X NP40 buffer was prepared. Lysates were pelleted on a magnetic stand at 4°C and supernatant was discarded. 300 µl of 1X NP40 buffer was added to each sample and samples were resuspended by nutation at 4°C. Once samples were thoroughly resuspended, they were pelleted on a magnetic stand. These washing steps were repeated 6 times. Following the final washing steps, beads were resuspended in 25 ul of 1X NP40 Buffer. 5 µl of beads were set aside for Western and the remaining beads were stored at −80°C in 100 µl of Trizol. SDS loading buffer was added was added to a 1X working concentration and the sample was heated at 95°C for 5m and stored at −20°C or used for Western (refer to Western Blot section).

### Helicase RNA IPseq

RNA was purified as previously described. RNA yield was quantified using Qubit or nanodrop according to manufactures instructions. RNA was run on a Fragment Analyzer according to manufactures instructions to assess quality. Inputs were diluted 1:50 to bring them into a similar range as the IgG and IP samples. To each sample 0.5 ng of Promega Luciferase Control RNA was added as a spike-in. Libraries were prepared as previously described except Poly(A) selection steps were skipped and library preparation was started with between 1-100 ng of total RNA. Reads were mapped to the M21017.1 NCBI *Drosophila* rRNA sequence record and the sequence of Luciferase obtained from Promega. All further analysis was performed using custom R scripts. Reads were assigned to features using featureCounts based off of a custom GTF file assembled based off of the Flybase record of rRNA sequences. Reads mapping to rRNA were normalized to reads mapping to the Luciferase spike-in control. Reads were further normalized to the reads from the corresponding input library to account for differences in input rRNA concentration between replicates and replicates were subsequently averaged. Tracks were visualized using the R package ‘ggplot’, with additional formatting performed using ‘scales’ and ‘egg’. The rRNA GTF was read into R using ‘rtracklayer’ and visualized using ‘gggenes’. Average reads mapping to rRNA from IgG control and IP was plotted and a one-sided bootstrapped paired t-test for was performed on regions on rRNA that appeared to be enriched in the IP samples compared to the IgG control as it is a non-parametic test suitable for use with low n using R with 100,000 iterations.

### Larp Gel Shifts

#### Cloning, Protein expression and purification

The Larp-DM15 protein expression construct (amino acids 1330-1481 corresponding to isoform D) was cloned into a modified pET28a vector by PCR using cDNA corresponding to accession ID NP_733244.5. The resulting fusion protein has an N-fHis_10_-maltose binding protein (MBP)-tobacco etch virus (TEV) protease recognition site tag. Protein expression and purification were performed as described previously (Lahr et al., 2015). Briefly, plasmid was transformed into BL21(DE3) *E. coli* cells and plated onto kanamycin-supplemented agar plates. A confluent plate was used to inoculate 500 mL of autoinduction media (Studier, 2005). Cells were grown for three hours at 37°C and induced overnight at 18°C. Cells were harvested, flash frozen, and stored at −80°C.

Cells were resuspended in lysis buffer (50 mM Tris, pH 8, 400 mM NaCl, 10 mM imidazole, 10% glycerol) supplemented with aprotinin (Gold Bio), leupeptin (RPI Research), and PMSF (Sigma) protease inhibitors. Cells were lysed via homogenization. Lysate was clarified by centrifugation and incubated with Ni-NTA resin (ThermoScientific) for batch purification. Resin was washed with lysis buffer supplemented with 35 mM imidazole to remove non-specific interactions. His_10_-MBP-DM15 was eluted with 250 mM imidazole. The tag was removed via proteolysis using TEV protease and simultaneously dialyzed overnight (3 mg TEV to 40 mL protein elution). Larp-DM15 was further purified by tandem anion (GE HiTrap Q) and cation exchange (GE HiTrap SP) chromatography using an AKTA Pure (GE) to remove nucleic acid and protein contaminants. The columns were washed with in buffer containing 50 mM Tris, pH 7, 175 mM NaCl, 0.5 mM EDTA, and 10% glycerol and eluted with a gradient of the same buffer containing higher salt (1 M NaCl). Fractions containing Larp-DM15 were pooled, and 3 M ammonium sulfate was added to a final concentration of 1 M. A butyl column (GE HiTrap Butyl HP) was run to remove TEV contamination. The wash buffer contained 50 mM Tris, pH 7, 1 M ammonium sulfate, and 5% glycerol, and the elution buffer contained 50 mM Tris pH 7 and 2 mM DTT. Fractions containing Larp-DM15 were buffer exchanged into storage buffer (50 mM Tris pH, 7.5, 250 mM NaCl, 2 mM DTT, 25% glycerol), flash frozen in liquid nitrogen, and stored at −80°C. The purification scheme and buffer conditions were the same as with *Hs*DM15 (Lahr et al., 2015), except cation and anion exchange buffers were at pH 7, as noted above.

### RNA preparation

5’-triphosphorylated *RpL30* and *Non1* 42-mers were synthesized (ChemGenes). Purine-substituted controls were synthesized by *in vitro* transcription using homemade P266L T7 RNAP polymerase (Guillerez et al., 2005). The transcription reaction containing 40 mM Tris, pH 8, 10 mM DTT, 5 mM spermidine, 2 mM NTPs, and 10-15 mM MgCl_2_ was incubated at 37°C for 4 hours. Transcripts were subsequently purified from an 8% polyacrylamide/6M urea/1XTBE denaturing gel, eluted passively using 10 mM sodium cacodylate, pH 6.5, and concentrated using spin concentrators (Millipore Amicon). All oligos were radioactively capped using Vaccinia virus capping system (NEB) and [α-^32^P]-GTP (Perkin-Elmer). Labelled oligos were purified using a 10% polyacrylamide/6M urea/1XTBE denaturing gel, eluted with 10 mM sodium cacodylate, pH 6.5, and concentrated by ethanol precipitation.

The RNA sequences used were:

RpL30: <u>CUUUU</u>GCCAUUGUCAGCCGACGAAGUGCUUUAACCCAAACUA Non1: <u>CUUUUU</u>GGAAUACGAAGCUGACACCGCGUGGUGUUUUUGCUU *Purine-substituted RPL30 control: <u>GAAAAG</u>CCAUUGUCAGCCGACGAAGUGCUUUAACCCAAACUA *Purine-substituted Non1 control: <u>GAAAAAG</u>GAAUACGAAGCUGACACCGCGUGGUGUUUUUGCUU

Oligos used for run-off transcription

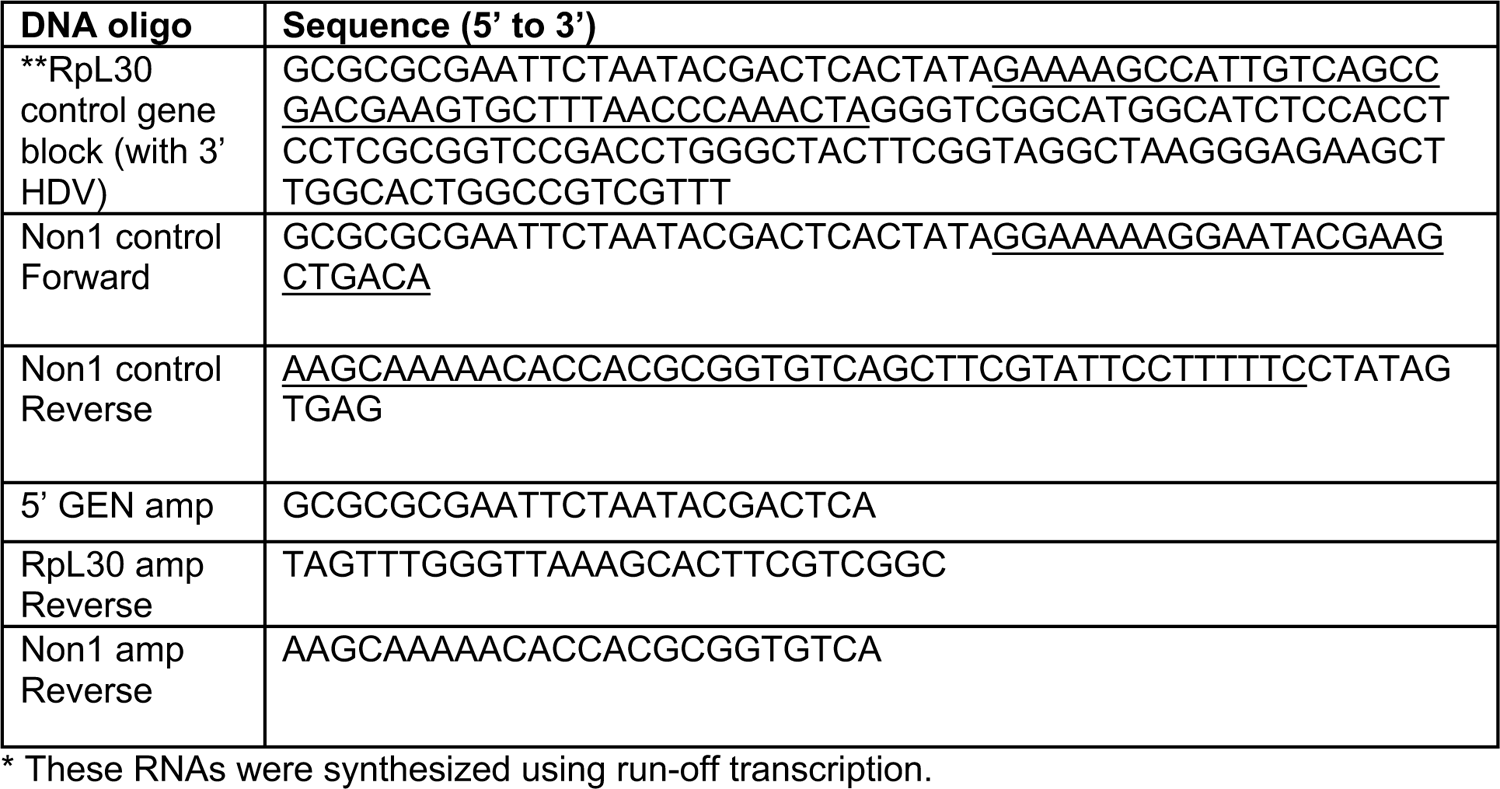

### Electrophoretic mobility shift assays (EMSAs)

Each binding reaction contained 125 total radioactive counts with final reaction conditions of: 20 mM Tris-HCl, pH 8, 150 mM NaCl, 10% glycerol, 1 mM DTT, 0.5 μg tRNA (Ambion), 1 μg BSA (Invitrogen), and <90 pM RNA. To anneal RNA, oligos were snap-cooled by heating at 95°C for 1 min and cooled on ice for 1 hour. For capped RpL30 shifts and capped purine-substituted controls, final concentrations of 0, 0.001, 0.003, 0.01, 0.03, 0.1, 0.3, 1, 3, 10, 30, and 100 nM Larp-DM15 were titrated. For capped Non1 shifts, final concentrations of 0, 0.01, 0.03, 0.1, 0.3, 1, 3, 10, 30, 100, 300, and 1000 nM Larp-DM15 were titrated. Native 7% polyacrylamide 0.5X TBE gels were pre-run on ice at 120 V for 30 min. Binding reactions were run at 120 V on ice for 45-52 min. Gels were dried for 30 min and allowed to expose overnight using a phosphor screen (GE). Screens were imaged using GE Amersham Typhoon. Bands were quantified using ImageQuant TL (GE). Background subtraction was first done using the rolling ball method and then subtracting the signal from the zero-protein lane from each of the shifted bands. Fraction shifted was determined by dividing the background-corrected intensity of the shifted band by total intensity of bands in each lane. Three independent experiments were done for each oligo, with the average plotted and standard deviation shown.

### Larp RNA IPseq

Larp IP was performed as described in the RNA IP-seq section above in triplicate. mRNA libraries were prepared as described in mRNAseq Library Preparation and Data Processing using a constant volume of RNA from each sample with input samples having been diluted 1:50. Data was processed as described as in the mRNAseq Library Preparation and Data Processing section. Targets are defined as genes with >2 fold enrichment and an adjusted p-value <0.05 in the Larp-IP libraries compared to input libraries, but not meeting those criteria in the IgG libraries compared to input.

### Larp RNA IP qPCR

Larp RNA IP was performed as described in the Larp RNA IPseq section with the following modifications. As the ovaries used were small, they were flash frozen in order to accumulate 40-50 ovaries for each biological replicate. Additionally, 5% input was taken for both RNA and protein samples. Once RNA was purified all of the RNA was treated with Turbo DNAse as in the mRNAseq Library Preparation and Analysis section. Reverse transcription (RT) was performed using Superscript II according to the manufacture’s protocol with equivalent volumes of RNA for each sample. cDNA was diluted 1:8 before performing qPCR using Syber Green. Each reaction consisted of 5ul Syber Green master mix, 0.4 ul water, 0.3 ul of each primer, and 4 ul of diluted cDNA. For each sample 3 biological and 3 technical replicates were performed. Oulier values of technical replicates were removed using a Dixon test with a cutoff of p<0.05. Remaining technical replicates were averaged, and the IP Input Ct value, the log_2_ of the Input dilution (20) was also subtracted to account for the Input being 5% of the total sample as follows:

ΔCt[normalized IP] = (Average Ct[IP] − (Average Ct[Input] − log_2_(Input Dilution Factor)))

Next, RNA recovery was normalized using the spike-in control for each sample as follows: ΔΔCt= ΔCt[normalized IP] − ΔCt[Luciferase]

Next, Each sample was normalized to it’s matched *bam* RNAi control as follows: *bam* RNAi normalized Ct= ΔΔCt[*aramis* RNAi IP] − ΔΔCt[[*bam* RNAi IP]

Finally, fold increase of IP from *aramis* RNAi over *bam* RNAi was calculated as follows: Fold Enrichment = 2^(−bam RNAi normalized Ct)^

Fold enrichment was plotted and One-sample t-test performed on *aramis* RNAi samples in R using a mu of 1.

